# Modeling Reveals How Direct-Acting Antivirals Redirect HBV Capsid Assembly Pathways to Noninfectious Products

**DOI:** 10.64898/2026.05.25.727729

**Authors:** Layne B. Frechette, Smriti Pradhan, Farzaneh Mohajerani, Carolina Pérez-Segura, Jodi A. Hadden-Perilla, Adam Zlotnick, Michael F. Hagan

## Abstract

Hepatitis B virus (HBV) infections cause chronic liver disease, resulting in about one million deaths per year, and there is currently no cure. Recent work has shown that a class of small molecules called capsid assembly modulators (CAMs) is promising for treating HBV. CAMs bind to HBV capsid protein subunits and alter their assembly, leading to non-functional and malformed structures rather than functional, closed shells. However, the mechanisms by which CAMs alter capsid assembly pathways remain unclear. Here, we extend a recently-developed kinetic Monte Carlo (KMC) model for HBV capsid assembly to simulate how CAMs affect assembly. In the model, CAMs alter assembly by preferentially binding to interfaces between certain quasi-equivalent subunit conformations. Simulations of the model reproduce experimental assembly product distributions. By analyzing assembly trajectories, we clarify the roles of thermodynamics and kinetics in determining assembly products, identify assembly mechanisms, and predict the key intermediates that lead to either capsids or malformed structures. Our findings enhance our fundamental understanding of capsid assembly, help advance the development of CAMs as a treatment for HBV and, more broadly, inform efforts to direct self-assembly pathways toward specific products.

## I. INTRODUCTION

During the lifecycle of many viruses, protein subunits self-assemble into closed shells (capsids) that enclose the viral genome and other viral components. Many viruses require precisely-controlled shapes and sizes to be infectious. For example, about half of all viruses have icosahedral capsids [1]. Yet, many viruses also exhibit polymorphism, or a tendency to form icosahedral capsids with different sizes or alternative, non-icosahedral structures [2–11]. Experiments and simulations have shown that small molecule antivirals [12, 13], as well as changing solution conditions [12, 14–17] or co-assembly with a cargo [12, 14–19] can affect which of these structures assemble. Understanding the mechanisms that control such polymorphic assembly would advance our understanding of the fundamental principles of self-assembly, aid in the rational design of drugs that misdirect virus assembly, and guide the development of synthetic self-assembly systems with tunable morphologies.

Icosahedral virus capsids are geometrically characterized by their triangulation number *T*, which (typically) corresponds to the number of different ‘quasi-equivalent’ conformations needed to assemble the capsid [20–22]. A capsid with triangulation number *T* has 60*T* subunits, with 60 subunits in each of the *T* conformations.

Hepatitis B virus (HBV) is a particularly interesting and important example of viruses that exhibit polymorphism. HBV is a major cause of human disease. Although an effective vaccine exists, hundreds of millions of people still suffer from chronic HBV infections, and in 2022, 1.1 million people succumbed to complications of HBV, including cirrhosis and liver failure [23]. There is currently no cure for HBV, but a number of therapeutics are under development [24]. Among these, capsid assembly modulators (CAMs) — small molecules that bind to the capsid subunits and alter their assembly properties — are particularly promising [25, 26]. CAMs can be divided into two major classes: CAM-Es, which accelerate assembly to form capsids that are morphologically normal but contain no genetic material [27–29], and CAM-As, which both accelerate and misdirect assembly to yield ‘aberrant,’ malformed structures [30, 31]. In this paper, we focus on CAM-As (which we abbreviate to ‘CAMs’ throughout), although our modeling framework can easily incorporate CAM-Es. Of the various CAM-As, heteroaryldihydropyrimidines (HAPs) have especially striking effects on HBV assembly behavior [12, 30]. HBV capsids are naturally dimorphic — while the majority of capsids that form in host cells have *T* = 4 structures, approximately 5% have *T* = 3 structures [32–34]. In vitro assembly of capsids leads to similar polymorphism, although the relative abundances of these polymorphs depends on the salt concentration, with higher salt concentrations correlating with progressively stronger association energy [14, 15, 35]. Kondylis et al. [12] showed that HAP-TAMRA (a fluorophore-labeled HAP [36] that serves as a model to study broader HAP-type CAM-A behavior) abolishes *T* = 3 capsid assembly and promotes the assembly of malformed structures, including large, extended sheets and structures containing a mix of capsid-like and sheet-like motifs. Investigating three regimens of low (80 mM), moderate (300 mM), and high (1000 mM) ionic strength, they also found that the relative abundances of *T* = 4 capsid and malformed structures depend on the salt concentration. The factors that determine these assembly products, and the underlying assembly pathways, remain unclear.

Computational and theoretical models have yielded important insights into the dynamics and assembly of HBV capsids with and without CAMs. Atomistic simulations of whole assembled capsids have shown that HBV capsid subunits are remarkably flexible [37], and that bound CAMs can significantly distort the capsid geometry [38, 39]; a survey of experimental structures supports this prediction [40]. Coarse-grained simulations enabled studying assembly pathways in the absence of CAMs, identifying long-lived intermediate structures and ‘hub states’ at which *T* = 3 and *T* = 4 assembly pathways diverge. Theoretical models [41, 42] have explored the roles of curvature free energy in polymorphism. Coarse-grained simulations [35] and theory [42] have helped explain how salt concentration affects the ratio of *T* = 3 to *T* = 4 capsids in the absence of CAMs.

A recent study by Kra et al. [13] combined time-resolved SAXS experiments, theory, and coarse-grained simulations to study how CAMs impact HBV assembly. The authors studied two different CAMs, JNJ-632 and Bay 41–4109: the latter induces assembly of large, non-*T* = 4/*T* = 3 shells, and both accelerate capsid assembly. Their models of SAXS data suggested that CAM binding accelerates assembly by increasing the subunit-subunit binding free energy, consistent with previous experiments [31]. They also performed coarse-grained simulations in which an assembling capsid is represented as a growing elastic shell. By varying the shell’s radius of curvature and Föppl-von Kármán number (which sets the ratio of stretching to bending moduli), they identified regimes where the shell assembled into *T* = 4 capsids, *T* = 3 capsids, and elongated shapes resembling the structures seen in experiments with Bay 41–4109. Because the elongated shapes formed at high Föppl-von Kármán numbers, Kra et al. [13] proposed that Bay 41–4109 could induce aberrant structures by changing the elastic modulus of a growing capsid. However, since their simulations did not explicitly represent CAM binding, they could not investigate this proposal or other mechanisms by which CAMs alter assembly pathways.

In this work we use computational modeling to understand the effect of CAMs on assembly pathways and products. We extend a coarse-grained, kinetic Monte Carlo (KMC) model that some of us previously developed to study HBV capsid assembly in the absence of CAMs [35]. The model was parametrized using experimental and atomistic simulation data, enabling it to capture protein-protein binding affinities and the mechanical and dynamical properties of capsids. The simulations are highly computationally tractable, which enables harvesting many assembly trajectories and analyzing the distribution of assembly pathways and outcomes. In the absence of CAMs, the model correctly predicted HBV dimorphism, how the relative abundance of *T* = 3 and *T* = 4 depends on solution conditions, and mechanisms of error correction by which overgrown structures shed excess subunits and close their shells.

In the extended model, CAMs bind to the interface between HBV dimers with an affinity that depends on the capsid protein conformation, based on experimental data. The KMC simulations qualitatively reproduce experimental assembly size distributions in the presence and absence of CAMs across a range of ionic strengths. In particular, the results are consistent with the experimental observation that CAMs lead to aberrant capsid sizes at moderate salt conditions (300mM NaCl, moderate association energy), but that high salt (1000mM NaCl, strong association energy) ‘rescues’ assembly of native-sized capsids. Because our simulations resolve the assembly/disassembly of individual dimers and CAMs, our analysis reveals assembly mechanisms and how CAMs disrupt them. Our results suggest that CAMs misdirect assembly by stabilizing subunit conformations that are incompatible with closed *T* = 4 or *T* = 3 capsids, and instead promote the formation of large sheet-like structures, consistent with electron microscopy observations. Moreover, using Markov state model analysis, we identify key assembly intermediates, including ‘hub states’ at which pathways between different assembly products diverge. Importantly, these predictions can be tested by experiments. By revealing the mechanism of action of CAMs, the computational results provide critical information for improving existing CAMs or designing new ones.

## II. RESULTS

We use a coarse-grained model to investigate the effects of CAMs on HBV capsid assembly. We have extended the elastic shell growth model developed in Ref. [35, 43] for HBV assembly in the absence of CAMs. Fig. 1 illustrates the model and compares the structures it forms to atomistic protein structures. Capsid protein dimers (Fig. 1Ai), the basic assembly subunits, are represented as edges in a flexible triangular mesh (Fig. 1Ai,ii,iii). CAMs are represented as beads that bind to dimer-dimer interfaces (Fig. 1B). We use a kinetic Monte Carlo (KMC) algorithm to simulate assembly and CAM binding/unbinding. The mesh assembles into *T* = 4 capsids (Fig. 1C), *T* = 3 capsids (Fig. 1D), or malformed structures such as sheets (Fig. 1E). The relative abundances of these different structures depends strongly on both solution conditions and the presence/absence of CAMs, as we will now describe. Details of the model and the KMC algorithm are given in Section IV.

**FIG. 1.**
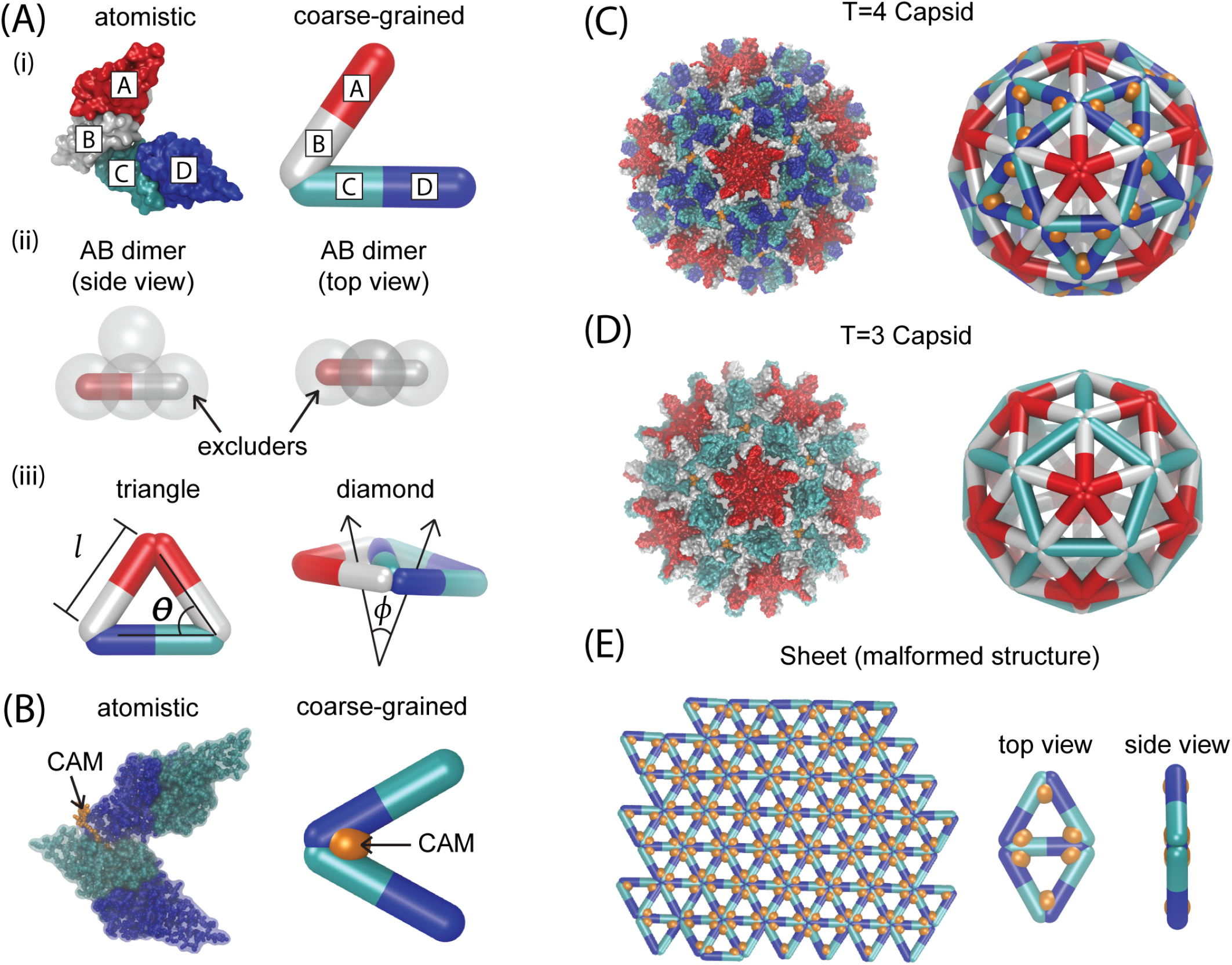
Overview of the coarse-grained model. **(A)** (i) Illustrations of atomic-resolution (left) and coarse-grained (right) dimer-of-dimers structures. The atomistic structure is extracted from the crystal structure of a complete capsid (PDB ID: 6BVF). Letters A, B, C, and D denote the four different quasi-equivalent conformations found in the *T* = 4 capsid. Here, an AB dimer (A in red, B in white) is bound to a CD dimer (C in cyan, D in blue). In the coarse-grained model, dimers are represented as edges in a flexible triangular mesh. (ii) Side (left) and top (right) views of a coarse-grained AB dimer. Transparent spheres are coarse-grained “excluder” particles. The excluders (of which there are four per dimer) are arranged so as to capture the overall dimer shape, with three excluders equally spaced along an edge and one placed above the middle of the edge (representing the “spike” in the atomistic dimer structure). For clarity, excluder particles are omitted from all other snapshots of coarse-grained structures. (iii) Illustration of a coarse-grained trimer-of-dimers or “triangle” (left) and a pentamer of dimers or “diamond” (right). The edge length *l*, binding angle *θ*, and dihedral angle *ϕ* all fluctuate about rest values which depend on the conformations of the dimers involved. **(B)** Left: atomic-resolution structure of a dimer of CD dimers, with a HAP-TAMRA molecule (an example of a capsid assembly modulator (CAM)) bound to the inter-dimer interface (PDB ID: 6BVF). Right: Coarse-grained dimer of CD dimers with a CAM (orange sphere) bound to the inter-dimer interface. **(C)** Illustration of the *T* = 4 capsid structure. Left: Atomistic structure of the *T* = 4 capsid, with HAP-TAMRA (orange) bound to the B-C and C-D interfaces (PDB ID: 6BVF). Right: Coarse-grained structure of the *T* = 4 capsid, with CAMs bound to the C-D interfaces. **(D)** Illustration of the *T* = 3 capsid structure. Left: Atomistic structure of the *T* = 3 capsid (PDB ID: 6BVN), with HAP-TAMRA (orange) bound to the B-C interfaces. Right: Coarse-grained structure of the *T* = 3 capsid. **(E)** Illustration of sheet-like malformed structures. Left: Example of a sheet from a simulation of the coarse-grained model. Right: Top and side views of a CAM-bound diamond of CD dimers. The rest dihedral angle for this structure is zero.

### A. Simulations capture how CAMs and salt concentration affect assembly products

Fig. 2 compares the distributions of assembly product sizes from our KMC simulations to experimental size distributions from Kondylis et al. [12]. We show results with and without CAMs at an ionic strength of *I* = 300mM, and with CAMs at higher (*I* = 1000mM) and lower (*I* = 80mM) salt concentrations. The simulations capture the key features observed in experiments: at *I* = 300mM without CAMs, the predominant assembly products have *n*_dimer_ ≈ 120, corresponding to *T* = 4 capsids. The simulations also exhibit a small peak at ≈ 90 dimers, corresponding to a *T* = 3 capsid; while *T* = 3 capsids have been observed in experiments at these conditions [12], the resistive-pulse experiments used for Fig. 2B could not resolve *T* = 3 from *T* = 4 capsids. Adding CAMs dramatically alters the size distribution – most assemblies have *n*_dimer_ > 120, and roughly half are nearly double that size (*n*_dimer_ ≥ 240, shown as a single peak). These large sizes correspond to malformed structures which fail to form a closed capsid. Instead, they contain extended, sheet-like motifs, sometimes with partially-formed capsid-like structures present at the periphery of the sheets. Increasing the ionic strength to *I* = 1000mM ‘rescues’ capsid assembly, partially restoring the peak at *n*_dimer_ = 120, although a significant number of assemblies still have *n*_dimer_ > 120. On the other hand, decreasing the ionic strength to *I* = 80mM abolishes the *n*_dimer_ = 120 peak and favors sheet-like assemblies, with most products having *n*_dimer_ ≥ 240. Notably, all these trends are qualitatively consistent between the KMC and experimental results.

**FIG. 2.**
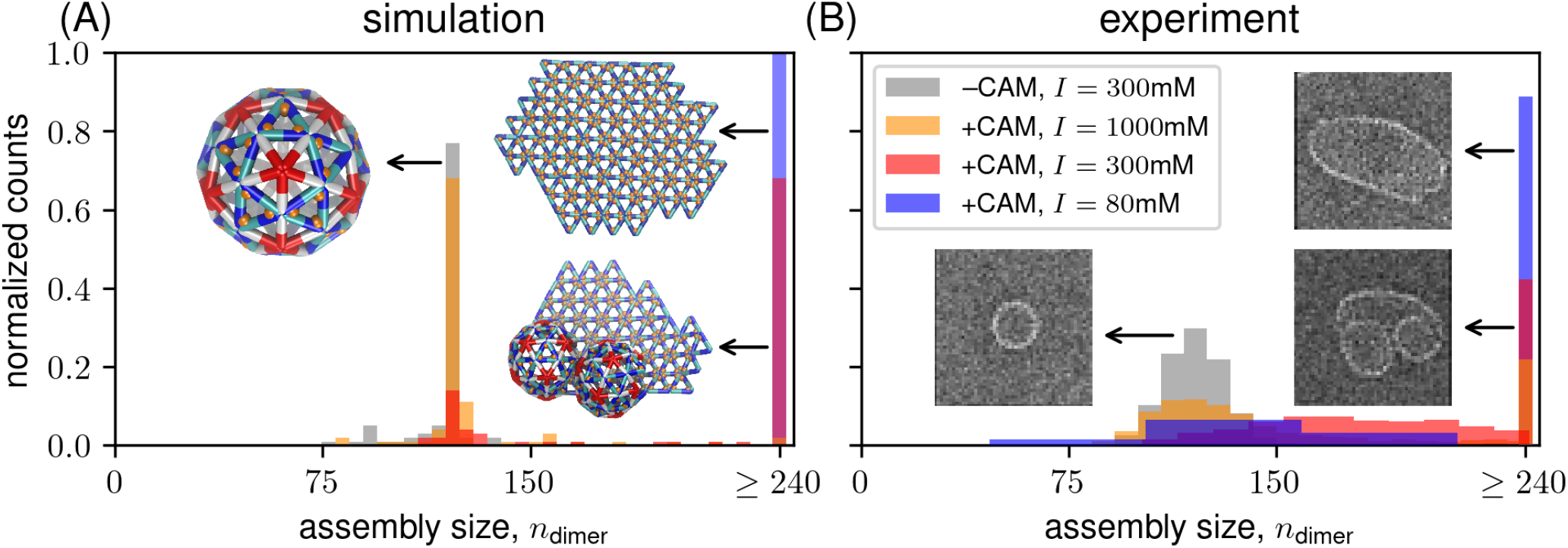
Simulations qualitatively reproduce experimental assembly product size distributions. Assembly product size distributions from **(A)** KMC simulations and **(B)** experiments [12] (data provided by S.C Jacobson). Results are shown for four different conditions: assembly without CAMs (–CAM) at an ionic strength of of *I* = 300mM, and assembly with CAMs (+CAM) at three different ionic strengths (1000, 300, and 80mM). The plots show the number of assemblies of different sizes (measured in number of dimers per complex, *n*_dimer_), normalized by the total number of assemblies for each condition. Insets show representative structures (simulation snapshots in panel A and electron micrographs in panel B, adapted from Ref. [12], copyright 2018 ACS Publications) for the different peaks. Malformed structures typically include extended sheets dominated by a hexagonal repeating structure, often with partially formed, *T* = 4–like shells at their periphery.

The simulations enable understanding the mechanisms underlying these experimental and computational observations. First, they can be partially understood from a thermodynamic perspective by considering how CAMs and salt concentration affect the free energies associated with dimer conformational changes and dimer-dimer binding. In our model, CAMs bind to C-D and D-C interfaces, stabilizing vertices with quasi-six-fold (hexameric) symmetry and hence favoring flat structures. This is consistent with the experimental size distributions in Fig. 2B and transmission electron micrographs showing flattening at quasi-six-fold axes in the presence of CAMs [12]. Increasing the salt concentration increases the total dimer-dimer binding affinity 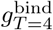 (Table I). Based on previous experimental work [44] and comparison between previous KMC results without CAMs and experimental size distributions in Mohajerani et al. [35], the trend in binding affinity primarily reflects the fact that increasing salt concentration decreases the difference in conformational free energy between AB and CD dimers 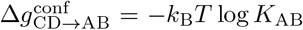, with 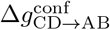 decreasing by 2*k*_B_*T* between *I* = 80mM and *I* = 1000mM (see Table I).

**TABLE I.** Simulation parameter values and descriptions.

| Salt-dependent interaction and thermodynamic parameters |  |  |
| --- | --- | --- |
| Parameter | Value(s) | Description |
| $\Delta g_{\text{CD} \rightarrow \text{AB}}^{\text{conf}}(k_B T)$ | $\begin{cases} 3.5, & I = 1000\text{mM} \\ 4.5, & I = 300\text{mM} \\ 5.5, & I = 80\text{mM} \end{cases}$ | conformational free energy difference between CD and AB dimers |
| $g_0^{\text{bind}}(k_B T)$ | $\begin{cases} -9.76, & I = 1000\text{mM} \\ -9.80, & I = 300\text{mM} \\ -9.85, & I = 80\text{mM} \end{cases}$ | reference dimer-dimer binding free energy |
| $g_{T=4}^{\text{bind}}(k_B T)$ | $\begin{cases} -9.62, & I = 1000\text{mM} \\ -9.42, & I = 300\text{mM} \\ -9.22, & I = 80\text{mM} \end{cases}$ | mean dimer-dimer binding free energy in $T = 4$ capsid |
| Relative interdimer binding affinities<br>(estimated from PDBePISA, Refs. [35, 84, 85]), with $g_{\text{D-C}}$ adjusted for assembly with CAMs) | | |
| Parameter | Value(s) | Description |
| $g_{\text{A-A}}/g_0^{\text{bind}}$ | 1.3 | A-A binding free energy |
| $g_{\text{B-C}}/g_0^{\text{bind}}$ | 1.1 | B-C binding free energy |
| $g_{\text{C-D}}/g_0^{\text{bind}}$ | 1 | C-D binding free energy |
| $g_{\text{D-B}}/g_0^{\text{bind}}$ | 0.9 | D-B binding free energy |
| $g_{\text{D-C}}/g_0^{\text{bind}}$ | 0.5 | D-C binding free energy |
| CAM interaction parameters |  |  |
| Parameter | Value(s) | Description |
| $g_{\text{C-D}}^{\text{CAM}}(k_B T)$ | -13 | binding free energy of CAM to DC-DC interface |
| $g_{\text{D-C}}^{\text{CAM}}(k_B T)$ | $-13 + (g_{\text{C-D}} - g_{\text{D-C}})$ | binding free energy of CAM to CD-CD interface |
| Chemical potentials and rate constants |  |  |
| Parameter | Value(s) | Description |
| $\mu_{\text{dimer}}(k_B T)$ | -11.5 | dimer chemical potential (corresponds to $\approx 10\mu\text{M}$) |
| $\mu_{\text{CAM}}(k_B T)$ | -10.1 | CAM chemical potential (corresponds to $\approx 40\mu\text{M}$) |
| $k_0^{\text{dimer}}(t_0^{-1})$ | $2 \times 10^{-2}$ | relative dimer binding rate<br>(ratio of attempted dimer binding moves to attempted vertex moves) |
| $k_0^{\text{CAM}}(t_0^{-1})$ | $2 \times 10^{-5}$ | relative CAM binding rate<br>(ratio of attempted CAM binding moves to attempted vertex moves) |
| Elastic and geometric parameters<br>(derived from all-atom simulations and capsid geometry, following Ref. [35]) |  |  |
| Parameter | Value(s) | Description |
| $\kappa_l(k_B T/\sigma^2)$ | 4200 | bond stretching modulus |
| $\kappa_\theta(k_B T)$ | 800 | binding angle bending modulus |
| $\kappa_\phi(k_B T)$ | 40 | dihedral angle bending modulus |
| $\sigma_i(\sigma)$ | 0.2 | excluder diameter |
| $l_{\text{AB}}^0(\sigma)$ | 0.95 | AB dimer rest length |
| $l_{\text{CD}}^0(\sigma)$ | 1.05 | CD dimer rest length |
| $\theta_{\text{A-A}}^0(\text{rad})$ | 1.17 | rest angle between BA and AB dimers (A-A interface) |
| $\theta_{\text{B-C}}^0(\text{rad})$ | 0.98 | rest angle between AB and CD(CC) dimers (B-C interface) |
| $\theta_{\text{D-B}}^0(\text{rad})$ | 0.98 | rest angle between CD(CC) and BA dimers (D-B interface) |
| $\theta_{\text{C-D}}^0(\text{rad})$ | 1.05 | rest angle between DC and DC dimers (C-D interface) |
| $\theta_{\text{D-C}}^0(\text{rad})$ | 1.05 | rest angle between CD and CD dimers (D-C interface) |
| $\phi_{\text{AB,capsid}}^0(\text{rad})$ | 0.48 | rest dihedral with AB interior edge in capsid |
| $\phi_{\text{CD,capsid}}^0(\text{rad})$ | 0.24 | rest dihedral with CD interior edge in capsid |
| $\phi_{\text{CD,malformed}}^0(\text{rad})$ | 0 | rest dihedral with CD interior edge in malformed structure (e.g. sheet) |

The thermodynamic factors discussed in the previous paragraph favor *T* = 4 capsid formation at *I* = 1000mM. However, our simulations show that thermodynamics alone cannot fully explain the rescue of capsids at *I* = 1000mM, i.e. why *T* = 4 capsids form instead of sheets. For example, if we artificially increase the CAM binding on- and off-rates while leaving the free energies unchanged with *I* = 1000mM, we observe a significant proportion of malformed, sheet-like structures (see Fig. S1). This observation underscores the importance that CAM binding be slow in comparison to dimer-dimer binding kinetics to observe the ‘rescue’ of *T* = 4 capsids under high salt, as previously suggested in Ref. [12].

Finally, we note that *T* = 3 capsids are absent with CAMs at *I* = 1000mM, although they make up a large fraction of assembly products *without* CAMs at *I* = 1000mM in both experiments and simulations (see Ref. [12] and Fig. S2). Our model suggests a thermodynamic rationale for this observation: CAMs stabilize C-D and D-C interfaces, which are not present in *T* = 3 capsids (see also Fig. 5 and the discussion of how CAMs destabilize *T* = 3 intermediates in Section III, Subsection B.2).

The observation that the low salt concentration (*I* = 80mM) favors sheets over *T* = 4 capsids also involves a combination of thermodynamic and kinetic effects. The increasing conformational free energy cost 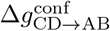 makes the the AB conformation, and thus icosahedral capsids, less favorable (reduces the magnitude of 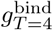, see Table I and SI Section S3), since the AB conformation is required for capsids but not sheets. From a kinetic perspective, destabilizing the AB dimers slows down *T* = 4 assembly, even in the absence of CAMs (see Fig. S3). Then, adding CAMs not only thermodynamically stabilizes C-D and D-C interfaces, but also promotes rapid nucleation of sheet-like structures (compared to slow nucleation of *T* = 4 capsids at low salt concentration without CAMs; see Fig. S3). Hence, sheets are also kinetically favored over *T* = 4 capsids at *I* = 80mM.

### B. CAMs alter assembly pathways

Next, to mechanistically understand the effect of CAMs on assembly products, we investigate how CAMs change assembly pathways. Fig. 3 shows representative trajectories for different salt concentrations leading to either *T* = 4 capsids or malformed structures (including sheets). We also provide movies of the assembly trajectories corresponding to these plots (Supplementary Movies S1-S5), see SI Section S5 for descriptions. Each panel in Fig. 3 shows the size (in number of dimers, *n*_dimer_) as a function of time for a single trajectory. In all cases, assembly is preceded by a long nucleation process in which the size fluctuates about the initial *n*_dimer_ = 3 nucleus. We show this phase explicitly in the top row (*I* = 1000mM); in the bottom two rows this waiting period is typically much longer, and so we start the plots at times near the beginning of the growth phase. We see that trajectories leading to *T* = 4 capsids proceed via a series of transiently stable intermediates which contain successively larger numbers of five-fold vertices (see Supplementary Movie S1). The same mechanism leads to *T* = 4 capsids at *I* = 1000 and 300mM (see Supplementary Movies S1 and S3), while no *T* = 4 capsids assembled for *I* = 80mM. By comparison to trajectories without CAMs (Ref. [35], Fig. S4, and Supplementary Movie S6), we see that the primary effect of CAMs on the *T* = 4 trajectories is to stabilize C-D interfaces (and hence CD conformations) in the three-dimer nucleus and subsequent small oligomers. This alters the structure of the first stable intermediate with *n*_dimer_ > 3: rather than the 10-dimer intermediate (which we henceforth refer to as the ‘decamer’) commonly observed in CAM-free capsid assembly (see Fig. 4B, Fig. 5, and Ref. [45]), we predominantly observe a 12-dimer intermediate (which we henceforth refer to as the dodecamer, see e.g. Fig. 5) in trajectories with CAMs. The dodecamer consists of a five-fold vertex with a CAM-bound dimer-of-CD-dimers. This structure is incompatible with a *T* = 3 capsid (in which adjacent triangles associated with five-fold vertices share a CC dimer edge). Thus, CAMs disfavor *T* = 3 capsid assembly by destabilizing the decamer, which is compatible with both *T* = 3 and *T* = 4 capsids, in favor of the dodecamer, which is compatible only with *T* = 4 capsids (see Section II, Subsection B.1 for further discussion).

**FIG. 3.**
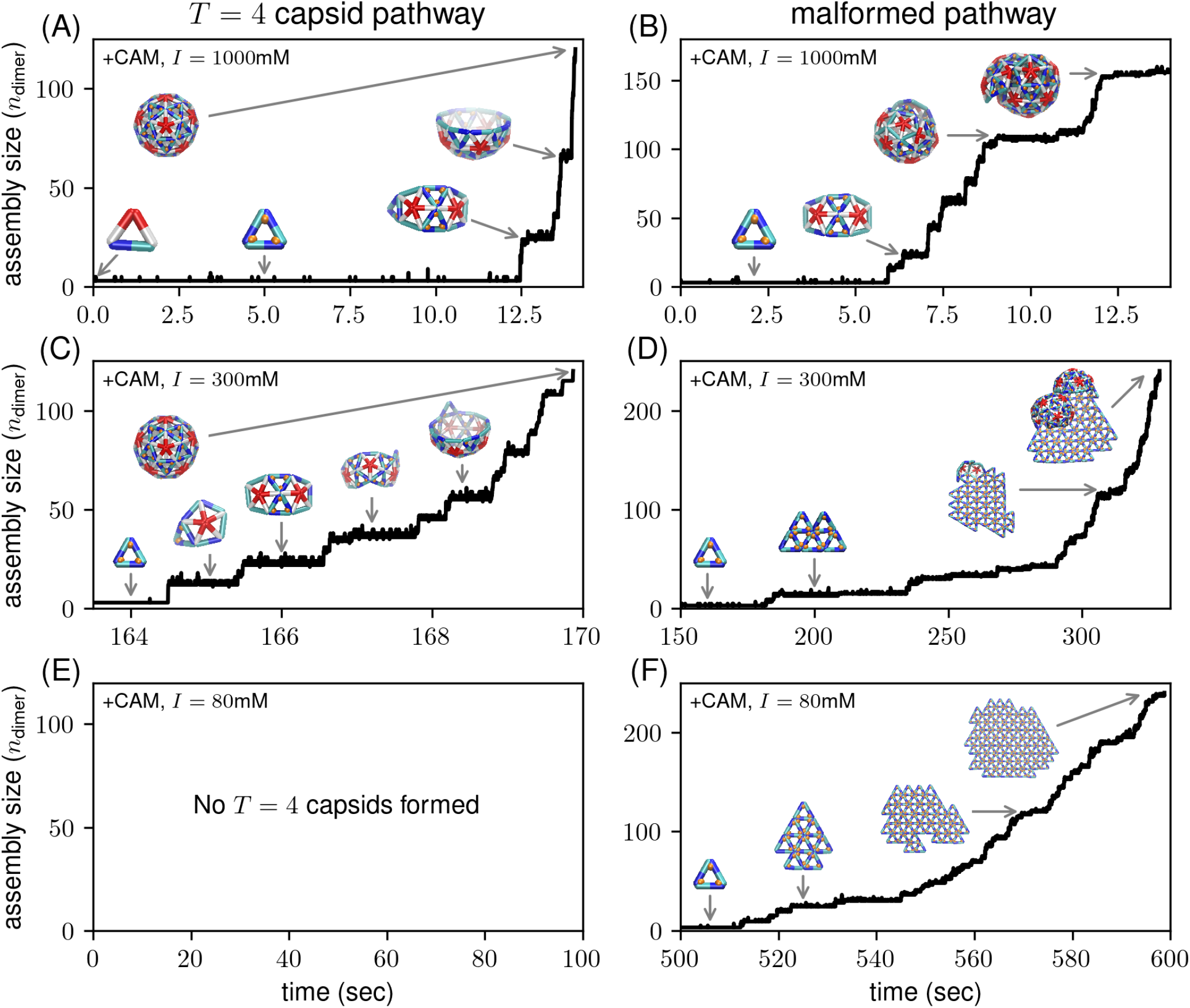
Changing salt concentration leads to different assembly pathways. The three rows show representative trajectories (assembly sizes versus time) leading to *T* = 4 capsids (left column, panels A, C, E) and malformed structures (right column, panels B, D, F) at different salt concentrations. CAMs are present in all cases. Representative intermediate structures are shown as inset snapshots with arrows pointing to the corresponding time points on the trajectories. See Supplementary Movies S1-S5 for movies of the assembly trajectories shown in panels A, B, C, D, and F.

**FIG. 4.**
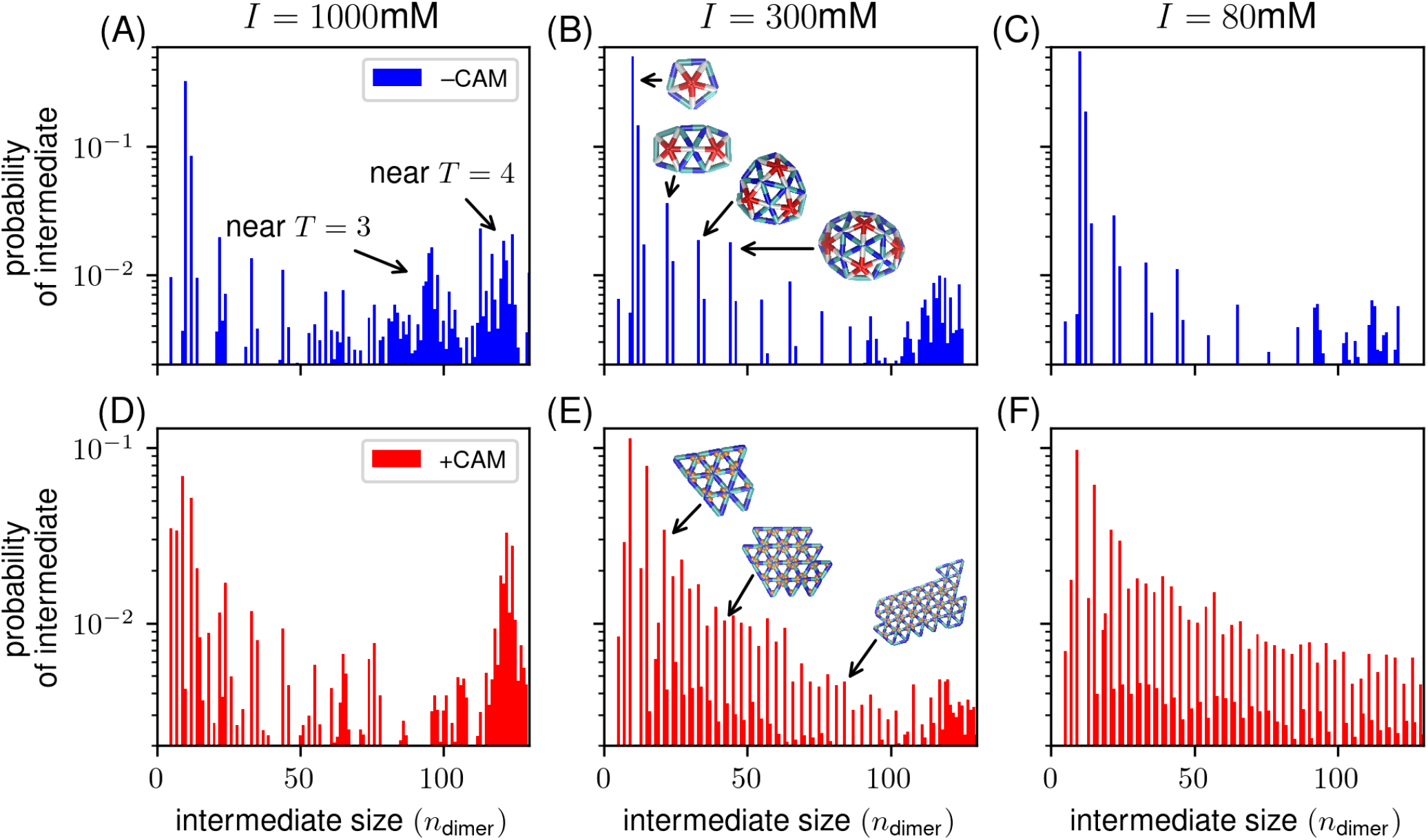
CAMs alter the prevalence of assembly intermediates. Probabilities of different assembly intermediates, normalized by the total number of intermediate structures (structures with size *n*_dimer_ *>* 3 which are neither *T* = 3 nor *T* = 4 capsids) over all trajectories at a given salt concentration and presence/absence of CAMs. The top row (panels A-C) shows results without CAM (–CAM, blue); the bottom row (panels D-F) shows results with CAM (+CAM, red). The three columns show results at different salt concentrations. Peaks corresponding to near-*T* = 3 and near-*T* = 4 capsid intermediates are labeled in panel A. In the middle column (panels B and E), snapshots illustrate representative structures associated with selected peaks in the distributions.

In contrast to *T* = 4 capsids, both the final structures and pathways associated with malformed assemblies change with salt concentration. At *I* = 1000mM, the predominant malformed structures are similar in size to *T* = 4 capsids, but fail to close. This is usually due to mixed *T* = 3/T = 4 morphologies (e.g. the final structure illustrated in Fig. 3B; see Supplementary Movie S2), which contain differing local curvatures that are incompatible with a closed shell. This class of pathways is very similar to the mixed-morphology malformed pathways we previously observed for assembly without CAMs [35]. However, we also observe other classes of malformed structures, including ‘dumbells’ consisting of two bound, partially-formed *T* = 4 capsids, as well as mixed *T* = 4/small sheet morphologies (see SI Fig. S5). At *I* = 300mM, most malformed structures consist of extended sheets, often decorated at their edges with partial, *T* = 4-like shells (final structure in Fig. 3D and Supplementary Movie S4). This suggests that the driving forces for sheet and *T* = 4 formation are comparable in magnitude under these conditions. In contrast, for *I* = 80mM, sheets become significantly more favorable (see Supplementary Movie S5). We rarely observe *T* = 4-like partial shells, and sheet growth is generally faster than at *I* = 300mM. While small (*n*_dimer_ ≲ 25) sheet-like intermediates grow via distinct jumps between transiently stable intermediates, larger assemblies grow more steadily. This is because the number of potential sites for adding additional CD dimers grows with the perimeter of the sheet: since the sheet is flat (rather than curved, as for *T* = 4 capsids) the perimeter increases steadily with the size of the sheet. The large number of available binding sites suggests that multiple sheet-like intermediates could associate to form larger malformed structures [46, 47], although we neglect this possibility in our model.

#### 1. CAMs alter the prevalence of key assembly intermediates

To better understand how CAMs modify and misdirect assembly pathways, we compute the probabilities of observing different intermediate structures during assembly. We define an intermediate as any structure with *n*_dimer_ > 3 which is not a complete capsid. Fig. 4 shows the probability distributions of intermediates as a function of size at different salt concentrations, both without (–CAM, blue, top row) and with (+CAM, red, bottom row) CAMs. Each distribution exhibits a series of peaks with a ‘multiplet’ structure, consisting of clusters of closely spaced peaks. For assembly without CAMs (top row), most multiplets are separated from their neighbors at a regular interval of ≈ 11 dimers. These correspond to intermediates with successively larger numbers of fivefold vertices. The multiplet structure arises due to a transient loss or gain of one or two dimers from the most probable structure in each multiplet. This is consistent with the step-like growth observed in the trajectories in Fig. 3. Dense clusters of peaks occur around *n*_dimer_ = 90 and 120, corresponding to near-*T* = 3 or near-*T* = 4 structures, respectively, awaiting the addition or removal of a few dimers to form a complete shell.

With CAMs, at *I* = 1000mM we observe a similar multiplet structure as in the trajectories without CAMs, although there are more numerous smaller peaks (likely due to the addition of dimers-of-CD-dimers, stabilized by CAMs, to structures with successively larger numbers of five-fold vertices), and the near-*T* = 3 peak is markedly reduced (consistent with the absence of *T* = 3 capsids in the *I* = 1000mM, +CAM histogram in Fig. 2).

At lower salt concentrations with CAMs, the intermediate probability distributions change dramatically. The near-*T* = 4 peak is markedly reduced at *I* = 300mM, reflecting the low yields of *T* = 4 capsids, and is entirely absent at *I* = 80mM, reflecting a total lack of *T* = 4 capsids. Moreover, the multiplet spacing changes from n_dimer_ = 11 to n_dimer_ = 3, corresponding to the addition of triangles of CD dimers rather than additional five-fold vertices. Fig. 4B,E show snapshots corresponding to the peaks at *I* = 300mM with and without CAMs, illustrating the structures of these intermediates. The intermediate probability distributions thus reflect the pathways by which different structures assemble. Importantly, the size distributions of assembly intermediates can be measured experimentally (e.g. [45, 48–51]), enabling a direct test of our predictions.

#### 2. Intermediate distributions reveal the mechanism by which CAMs suppress *T* = 3 capsid assembly

A closer inspection of small intermediate probabilities reveals the mechanism by which CAMs abolish *T* = 3 assembly. Fig. 5 shows the probability distributions of intermediates up to size n_dimer_ = 15 for assembly with and without CAMs at *I* = 1000mM. To facilitate comparison, we show each distribution normalized by the probability of the most likely intermediate for that system (thus, e.g. the probabilities for assembly without CAMs are normalized by the probability of the decamer). We show snapshots illustrating the most common structure at each size above the corresponding bars in the plot. We see that without CAMs, the most probable intermediate is the decamer, followed by the dodecamer, both of which have a single five-fold vertex. By contrast, the most probable intermediate with CAMs is a 9-mer consisting entirely of CD dimers (smaller all-CD 5-mer and 7-mer intermediates also feature prominently). The next most common intermediate is the dodecamer, and a 14-mer containing a single five-fold vertex with two additional dimers-of-CD-dimers is also common. Importantly, the probability of observing the decamer in trajectories with CAMs is tiny compared to the probability of the dodecamer and of the 14-mer. This observation provides strong evidence for the mechanism of abolishing *T* = 3 capsid assembly that was suggested by assembly trajectories (Fig. 3): the decamer is compatible with both *T* = 3 and *T* = 4 capsids, but due to their CD dimers, the dodecamer and the 14-mer are compatible only with a *T* = 4 capsid. Thus, by stabilizing the dodecamer and the 14-mer, CAMs greatly reduce the probability of forming a *T* = 3 capsid (or any other 90-mers, see Fig. 4A,D).

**FIG. 5.**
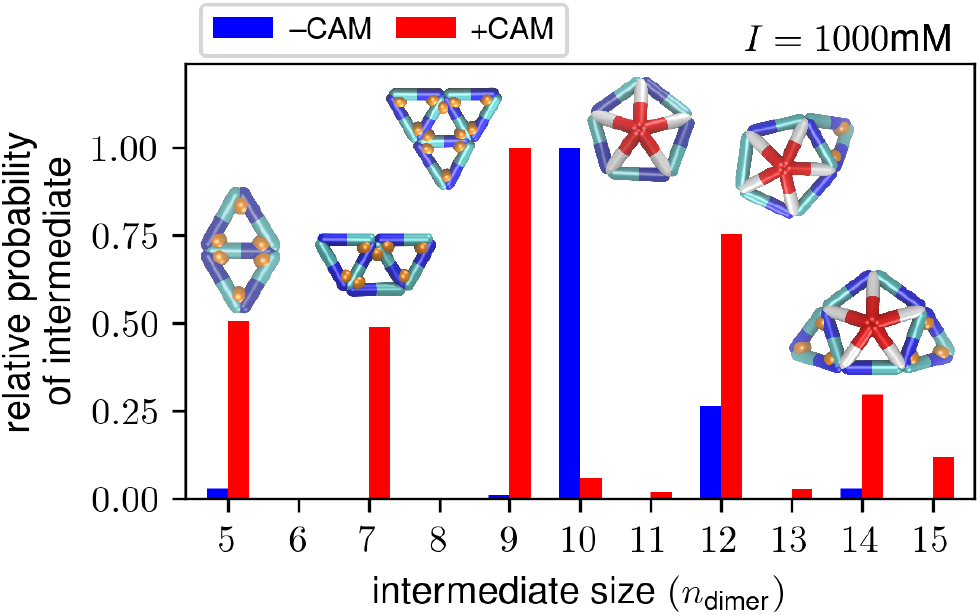
CAMs abolish *T* = 3 capsid formation by altering the distribution of early assembly intermediate sizes. Probability of different assembly intermediates (*n*_dimer_ *>* 3, excluding *T* = 4 or *T* = 3 capsids), relative to the probability of the most probable intermediate, as a function of intermediate size (measured in number of dimers, *n*_dimer_). We compute probabilities by counting the number of assemblies of each size within all trajectories at a given set of conditions. We show results for simulations at *I* = 1000mM without (blue) and with (red) CAMs. Snapshots illustrate representative intermediates corresponding to each peak.

#### 3. Intermediate distributions resolved by pathway identify small intermediates unique to different assembly pathways

The distributions in Fig. 4 illustrate the mechanism by which *T* = 4 capsids and sheets grow, but at what point does an assembly ‘choose’ one outcome over the other? This question is particularly relevant to assembly with CAMs at *I* = 300mM, where both *T* = 4 capsids and sheets form in significant numbers. To answer it, we partition trajectories into those that end in *T* = 4 capsids and those that end in sheet-like structures. We define a sheet-like structure as any structure which at least 10 ‘sheet-like CD dimers,’ in which the CD dimer edge is shared by two triangles, both of which consist solely of CD dimers (see e.g. Fig. 1E). (By contrast, ‘T = 4-like CD dimers’ form an edge that is shared by a triangle consisting of only CD dimers and a triangle with two AB dimers and one CD dimer; see Fig. 1Aiii, right.) This definition accounts for the fact that many malformed structures at *I* = 300mM contain both *T* = 4-like and sheet-like regions. We then compute intermediate probability distributions separately for trajectories ending in *T* = 4 capsids and trajectories ending in sheet-like structures (Fig. 6). *T* = 4 trajectories (turquoise) prominently feature the dodecamer and 14-mer, as well as a sheet-like 5-mer. In contrast, sheet-like pathways (orange) prominently feature the 9-mer, as well as the 7-mer and the larger 13-mer and 15-mer. Importantly, while the distributions exhibit some overlap for *n*_dimer_ ≤ 9, there is no overlap for *n*_dimer_ ≥ 12. Thus, sheet-like 13-mers almost never occur in *T* = 4 pathways, while dodecamers (with their five-fold vertices) almost never occur in sheet pathways. This suggests that the dodecamer is ‘committed’ to forming (i.e. most probably going to form) a *T* = 4 capsid (or malformed capsid) rather than a sheet, while the 13-mer is committed to forming sheet-like structures. Thus, assembly outcomes appear to be determined by which small intermediate – a *T* = 4-like dodecamer or a sheet-like 13-mer – forms first.

**FIG. 6.**
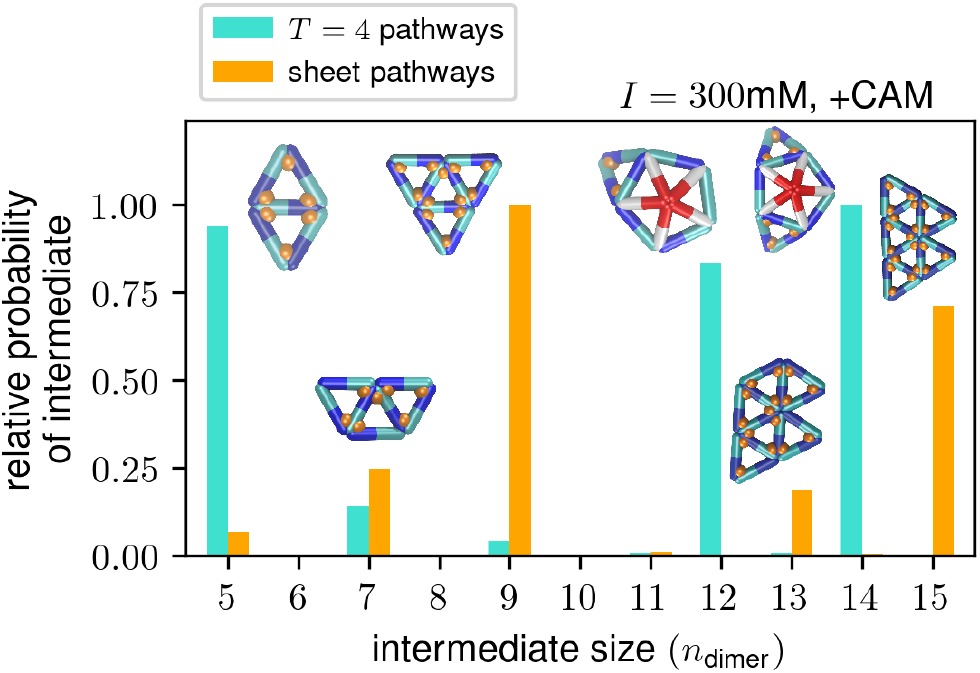
Small intermediates determine assembly products. Probability of different assembly intermediates (*n*_dimer_ *>* 3, excluding *T* = 4 or *T* = 3 capsids), relative to the probability of the most probable intermediate, as a function of intermediate size (measured in number of dimers, *n*_dimer_), for assembly with CAMs and *I* = 300mM. We compute probabilities by counting the number of assemblies of each size within all trajectories with CAMs at *I* =300mM. The plot shows intermediate probabilities for pathways that end in *T* = 4 capsids (turquoise) and sheets (orange). The insets show representative snapshots of intermediates corresponding to each peak.

#### 4. Committor probabilities identify small intermediate ‘hub states’ that determine assembly outcomes

To test whether different intermediates are indeed committed to forming certain structures, we computed committors for different intermediates. For a dynamical process involving transitions between a reactant state A and product state B, the committor probability, 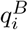, is the probability that a trajectory initiated in a state i reaches state B before state A [52, 53]. To compute committor probabilities, we first constructed a Markov state model (MSM) for HBV assembly with CAMs (see Section IV F for full details). An MSM coarse-grains a system’s dynamics into Markovian transitions between discrete microstates. We used a physically transparent discretization in which microstates 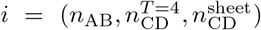 are characterized by the number of AB dimers (*n*_AB_), the number of *T* = 4-like CD dimers 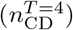, and the number of sheet-like CD dimers 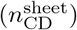 in an assembly. We used the transition matrix ***T*** (*τ*) obtained from the MSM to compute committor probabilities 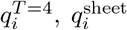 associated with reaching the product states (*T* = 4 capsid and sheet, respectively) before either returning to the reactant state (the initial, three-dimer nucleus) or ending in an alternative product state (e.g. a malformed capsid) (see Section IV G for details).

Plots of 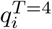 and 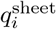 as a function of n_AB_, 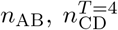 and 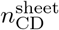 are shown in Fig. 7A,B. When 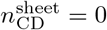 and 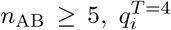 increases steadily with n_AB_ and 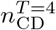 Note that while for most near-T = 4 states (with n_AB_ and 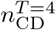 both close to 60) 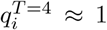, there are some states with 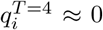, corresponding to malformed capsids. For nearly all microstates with 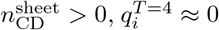. Conversely, 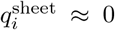 for 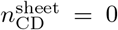, but steadily increases with increasing 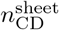. In particular, for the dodecamer, 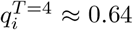 and 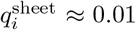; for the 13-mer, 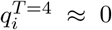 and 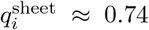. This confirms that the five-fold-vertex-containing dodecamer is indeed committed to forming a *T* = 4 capsid, while the sheet-like 13-mer is committed to forming a sheet. Additionally, we identify “hub states” as intermediates for which either 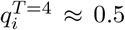 or 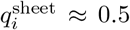, i.e., they are equally likely to either form the structure of interest (*T* = 4 capsid or sheet) or not. We show snapshots of representative hub states in Fig. 7C. The left state has 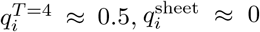, contains a single five-fold vertex, and has 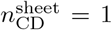. For the right state, 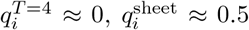; the structure has two five-fold vertices and a small sheet-like domain. Interestingly, there are no hub states with both 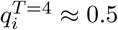 and 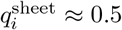, suggesting that trajectories must return to the initial three-dimer nucleus to switch between *T* = 4 and sheet assembly pathways. In Fig 7D we show representative hub states for assembly without CAMs, obtained by constructing an MSM from trajectories without CAMs at *I* = 300mM and identifying states with 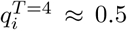 (see Fig. S6 for committor plots). In contrast to the hub states with CAMs shown in Fig. 7C, the hub states without CAMs exhibit mixed *T* = 4/T = 3 morphologies, consistent with the results of Mohajerani et al. [35]. Thus, without CAMs, assembly outcomes are determined by a competition between locally *T* = 3-like and *T* = 4-like morphologies, whereas with CAMs they are instead shaped by a competition between locally sheet-like and *T* = 4-like morphologies.

**FIG. 7.**
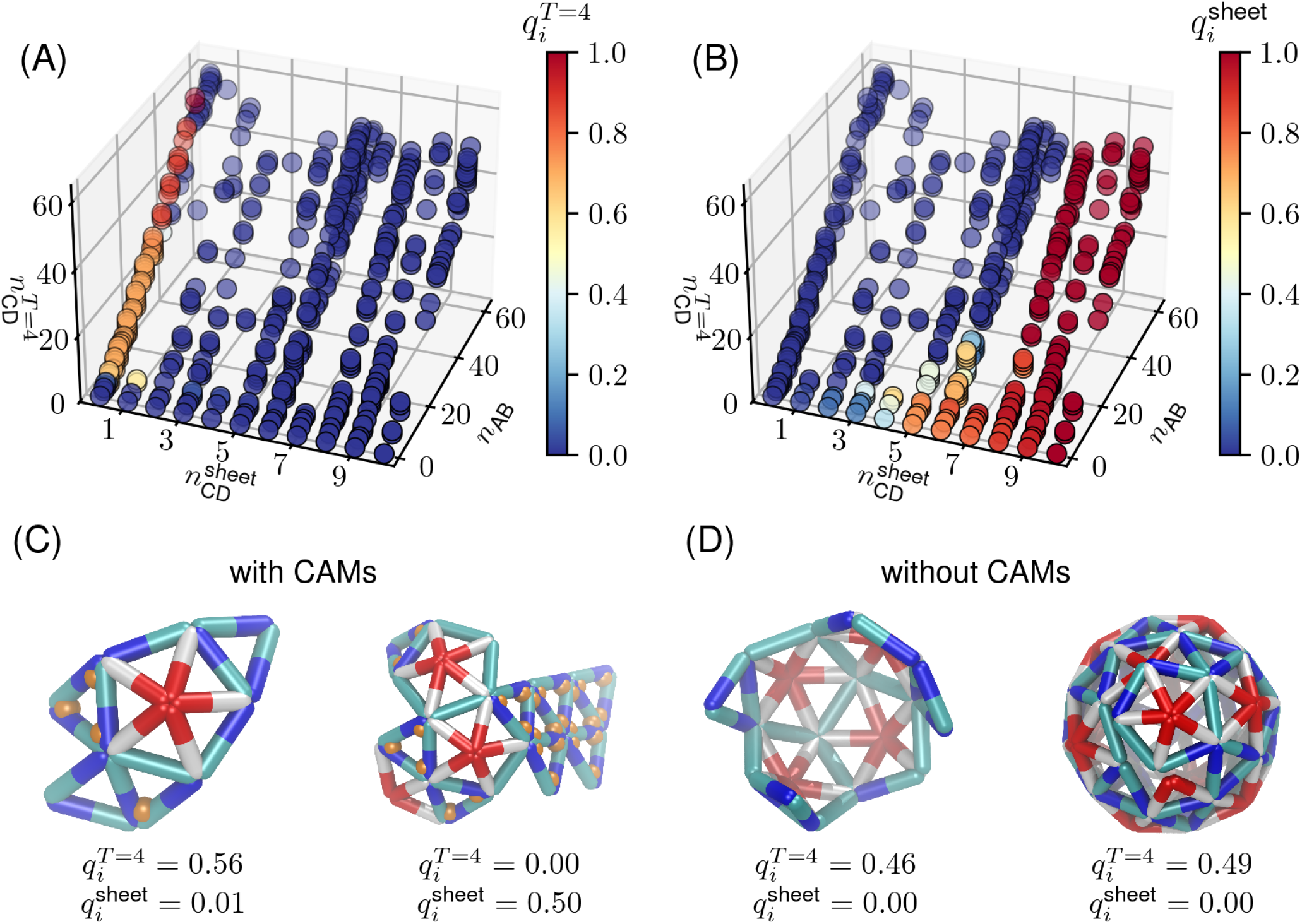
Committor probabilities enable identifying hub states for *T* = 4 and sheet assembly. (A) Committor probabilities 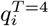 for *T* = 4 capsid assembly as a function of *n*_AB_, 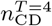, and 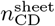. The color indicates the value of 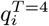. (B) Committor probabilities 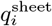 for sheet assembly (any structure with 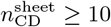 as a function of *n*_AB_, 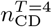, and 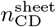. (C) Representative snapshots of *T* = 4 capsid hub states with 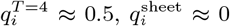 (left, 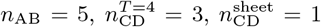) and sheet hub states with 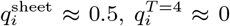 (right,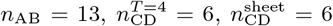). The committor probabilities are listed below the snapshots. In panels (A-C), there are CAMs and *I* = 300mM. (D) Representative snapshots of *T* = 4 capsid hub states without CAMs at *I* = 300mM. Both the left and right snapshots have 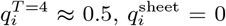 (sheets do not form without CAMs).

## III. DISCUSSION

We have presented a computational framework that, while parameterized from atomistic simulations, is sufficiently tractable to simulate assembly timescales. We have shown that the model predicts assembly kinetics and products consistent with experiments on the effect of small molecule antiviral agents (CAMs) on HBV assembly. The model results enable explaining how assembly products depend on salt concentrations and CAMs in terms of the thermodynamic affinities and kinetic binding rates of capsid protein dimers and CAMs, as well as the effects of CAM binding on the local geometry of a growing partial capsid. Moreover, the model makes experimentally testable predictions about how CAMs affect assembly pathways as a function of salt concentration, including the structures and concentrations of specific intermediates.

In this section, we further discuss how the model predictions can be experimentally tested, limitations of the current model and how they might be overcome, and potential applications to other assembly systems.

### A. Testing in experiments

While the predicted assembly product distributions are consistent with existing experiments (Fig. 2), our simulations also provide extensive predictions that could be tested by further experiments. These include assembly kinetics and predictions of assembly intermediate populations and lifetimes, which could be tested using high-resolution, time-resolved experimental techniques, such as time-resolved SAXS [45], resistive pulse sensing [48– 51], interferometric scattering [54–57], and mass photometry [58–60]. Consistency of these experimental measurements with our predictions would support our proposed mechanisms, including how CAMs suppress *T* = 3 capsid formation, and which intermediates are hub states, e.g. lead to *T* = 4 capsids versus sheets.

### B. Limitations of the model

We have made the simplifying assumptions that CAMs only bind to interfaces between CD dimers, and that the only effect of CAM binding is to increase the net dimer-dimer binding affinity. However, experimental evidence and all-atom simulations show that some CAMs (including HAPs) not only increase dimer-dimer affinities, but also distort dimer-dimer binding angles, favoring the flat angles found at quasi-sixfold axes. For example, HAP-TAMRA and HAP18 asymmetrically distort capsids, flattening CD conformations and leading to faceted structures [36, 39]. The binding of HAP-ALEX (another fluorophore-labeled HAP) to intact *T* = 4 capsids exhibits negative cooperativity [61], suggesting that distortions in binding geometries are correlated beyond neighboring subunits. Our model partially accounts for this effect by having CAMs bind only to the CD dimer conformations present at the quasi-sixfold axes. The fact that *g*_D-C_ needs to be carefully tuned within a range −4.88*k*_B_*T* to − 4.93*k*_B_*T* — small enough to destabilize *T* = 4 capsids under normal assembly conditions (without CAMs), but large enough to allow sheets to grow when CAMs are present, suggests that this parameter may be qualitatively accounting for more complex conformational changes during assembly.

The effects of CAMs on binding angles could be more completely incorporated by making the preferred dimer-dimer binding angle and/or dihedral angle change upon CAM binding. However, the effects of CAM binding on subunit angles in the atomistic simulations of complete HBV capsids are limited by constraints arising from the neighboring subunits, and the resulting changes in the coarse-grained binding angles are small. Moreover, the remarkable flexibility of HBV dimers [37, 38] suggests that they may sample alternative conformations when not part of a capsid. Thus, accurately parameterizing the effects of CAMs on binding angles will require data from all-atom simulations of smaller intermediates in which fewer constraints from the capsid geometry are present, and potentially adding additional degrees of freedom to the coarse-grained model. However, these steps are beyond the scope of the current work.

### C. Outlook

The computational framework used in the present article can be readily generalized to other antiviral molecules or viruses, provided that atomistic simulation data and experimental data on binding affinities and rates is available for parameterization. For example, while this work was motivated by experiments looking at the effect of a particular CAM, HAP-TAMRA, on assembly, other CAMs have different effects on assembly [26, 62]. Moreover, researchers are investigating other potential antiviral molecules that perturb capsid assembly for a wide variety of viruses (e.g. [63–67]).

More broadly, our results elucidate the engineering principles that natural viruses have evolved to optimize their assembly, and how assembly pathways can be redirected to alternative structures. These insights could advance efforts in the emerging field of “synthetic structural biology” [68] to design de novo subunits that efficiently self-assemble into large, complex, and functional structures. For example, our results show that the binding of effector molecules to assembly subunits could be a powerful strategy for manipulating assembly morphologies. This strategy could be implemented synthetically by designing multicomponent assembly systems, e.g. using DNA origami [69–72] or protein design [73–79], in which small DNA strands [80] or peptides [81, 82] bind preferentially to the interfaces between certain components, perhaps stabilizing local geometries consistent with different assembled structures (e.g. nanotubes with different curvatures) [83]. Such synthetic systems could provide an alternative, controllable avenue for understanding the principles by which effector molecules stabilizing certain subunit conformations can control self-assembly.

## IV. METHODS

### A. Model

Our coarse-grained model for HBV assembly with CAMs extends the elastic shell growth model developed in Ref. [35, 43] for HBV assembly in the absence of CAMs. As described in Section II, capsid protein dimers (subunits) are represented as edges in a flexible triangular mesh (Fig. 1Ai,ii,iii). We characterize the mesh geometry by its edge lengths l, binding angles θ between two bound edges, and dihedral angles ϕ between pairs of triangles that share an edge. Each edge can switch between two conformations — AB and CD/CC — where A, B, C, and D refer to the four quasi-equivalent monomer conformations appearing in *T* = 4 HBV capsids. The CD/CC designation refers to the fact that *T* = 4 capsids have AB and CD asymmetric dimers whereas *T* = 3 icosahedra have AB asymmetric dimers and CC symmetric dimers. When referring to a dimer-dimer interface, we only list the two monomers that are in contact; thus, e.g., the interface between a DC-DC dimer of dimers is referred to as the C-D interface. The lengths, binding angles, and dihedral angles fluctuate about their rest values, which depend on the conformations of the dimers involved (see Table I); deviations from the rest values incur an elastic energy cost.

In atomistic protein structures, CAMs such as HAP-TAMRA are bound to dimer-dimer interfaces (Fig. 1B, left). In our model, we represent CAMs as single beads that bind to the corners of the triangles (which represent dimer-dimer interfaces; Fig. 1B, right). Importantly, when two dimers bind, one acts as a “pocket” and the other acts as a “cap” [40]; thus, e.g., C-D and D-C interfaces are distinct. We represent this asymmetry in our model by using a directed half-edge data structure (see SI Section S2 A for details). Cryo-EM data indicate [36] that HAP-TAMRA – the CAM used in the experiments that inspired this work – primarily binds to quasi-sixfold vertices (i.e. B-C and C-D interfaces). However, the experiments of Kondylis et al. [12] suggest CAM binding favors flatter structures, and previous simulations by Mohajerani et al. suggest that extended, flat structures are composed primarily of dimers in the CD conformation. Motivated by these observations and the hypothesis that CAM binding to B-C interfaces is not essential for malformed assemblies (supported by atomistic simulations showing that B-C interfaces are “CAM-ready” and hence little prone to inducing structural distortions in capsids [39]), we reduce the number of parameters by considering a model in which CAMs bind only to only to C-D interfaces (found in *T* = 4 capsids) and D-C interfaces (present in sheet-like structures, but not *T* = 4 or *T* = 3 capsids). The model could be easily extended to allow for binding to other interfaces as well.

We use a kinetic Monte Carlo (KMC) algorithm to simulate the dynamics of the mesh. We work in the grand canonical ensemble, where chemical potentials *µ*_dimer_ and *µ*_CAM_ set the concentrations of dimers and CAMs, respectively, in a reservoir. Since the concentrations are constant in our simulations, we simulate dynamics over time periods in the reaction which are short in comparison to the depletion of subunits or CAMs. The concentrations of CD and AB dimers in the reservoir follow from mass conservation and by the noting that the conformational free energy difference determines the ratio of their concentrations in the reservoir: [AB]/[CD]=exp 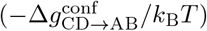. Starting from an initial, three-dimer nucleus (Fig. 1Aiii, left), assembly proceeds by the addition of edges or pairs of edges, which represent the binding of dimers or dimers-of-dimers to the growing structure. Simultaneously, CAMs can bind/unbind to/from the corners of the triangles. Importantly, dimer-dimer and CAM-dimer-interface binding are reversible, and the relative rates of binding and unbinding are set by the dimer-dimer and CAM-dimer-interface binding free energies. We describe the model parameters and the KMC algorithm in more detail in the following subsections and in the SI.

The *T* = 4 capsid corresponds to a free energy minimum in our model, which is the global minimum under most conditions that we simulate, except with CAMs at low salt concentration where sheets are favored (*I* = 80mM; see SI Section S3). Fig. 1C illustrates the atomistic and coarse-grained structures of the complete capsid. The atomistic structure has HAP-TAMRA bound to B-C and C-D interfaces; the coarse-grained structure has CAMs bound only to the C-D interfaces. The rest values of dihedral angles (i.e. the preferred angle between the two triangles in each diamond) are such that the *T* = 4 capsid minimizes the elastic energy. However, as shown by Mohajerani et al. [35], the diamonds are sufficiently flexible that *T* = 3 capsids can also assemble (Fig. 1D). Additionally, CAMs can stabilize diamonds that consist only of CD dimers (Fig. 1E, right). These diamonds contain both C-D interfaces (present in *T* = 4 capsids) and D-C interfaces (absent from *T* = 4 capsids), and have a rest dihedral angle of zero. This stabilizes flat, non-capsid (“malformed”) structures, which we refer to as “sheets” (Fig. 1E, left).

### B. Units

We set σ = 8nm as our unit of length, t_0_ = 10^−5^sec/sweep as our unit of time (where a sweep consists of a certain number of attempted Monte Carlo moves, as defined in Subsection C below), and *k*_B_*T* as our unit of energy, where k_B_ ≈ 1.381 × 10^−23^kg(m/s)^2^/K is Boltzmann’s constant and *T* = 300K is the temperature in Kelvin. We express all other quantities in terms of σ, t_0_ and *k*_B_*T*.

### C. Coarse-grained energy function

The KMC dynamics obey detailed balance with respect to a coarse-grained energy function:

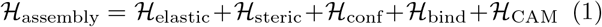

where the terms on the right-hand side account for, respectively, elastic, steric (excluded volume), conformational, dimer-dimer interaction, and dimer-CAM interaction free energies. The elastic energy function is:

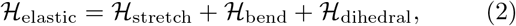

where the stretching, bending, and dihedral elastic energies are given respectively by:

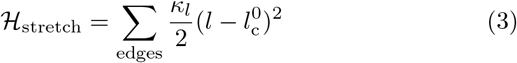

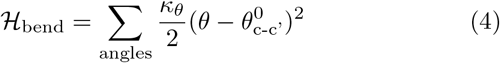

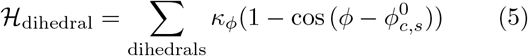

Here, l is the (instantaneous) length of an edge, θ is the angle between two edges within a triangle, and ϕ is the dihedral angle between two triangles; 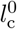 and 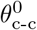 are the rest length and rest angle, whose values depend upon the conformations *c* of the edges involved (see Table I in the Appendix); and κ_*l*_ = 4200*k*_B_*T*/σ^2^, κ_*θ*_ = 800*k*_B_*T*, and κ_*ϕ*_ = 40*k*_B_*T* are elastic moduli, whose values are inferred from atomistic molecular dynamics simulations of intact capsids [35]. (Note that to simplify notation, we write e.g. 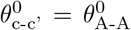 when *c* =BA, *c*′ =AB.) The quantity 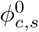 is the rest dihedral angle, which depends on the conformation *c* of the interior edge of the diamond that forms the dihedral and the conformations of the other edges in the diamond. Although the complete energy function depends on the conformations of all five dimers involved in a dihedral, we simplify the presentation in the text by writing s = capsid, malformed. Here “capsid” denotes diamonds that are present in *T* = 3 and *T* = 4 capsids and “malformed” refers to all other diamonds (which are found in mis-assembled structures). The most common malformed diamonds formed in our simulations are those that consist only of CD dimers (i.e. Fig. 1E).

The steric energy function, which enforces dimer excluded volume, is:

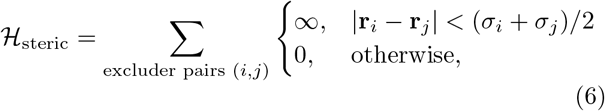

where σ_*i*_ is diameter of a hard sphere “excluder” pseudoatom. There are four excluder pseudoatoms per dimer (edge), with three evenly spaced along the edge and one above the edge, the latter representing the dimer “spike” (see Fig. 1Aii). We set σ_*i*_ = 0.2σ for all excluders i.

The conformational energy function ℋ_conf_ accounts for the intrinsic free energy difference between different quasi-equivalent conformations 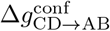, which is related to their relative concentrations at equilibrium by 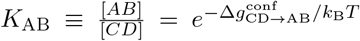. Given their high degree of structural similarity, we set the CD and CC conformations to have the same intrinsic free energy 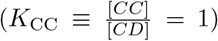, and we set the CD/CC dimer as the reference conformation with zero conformational free energy. Hence, only the conformational free energy of AB dimers relative to CD/CC dimers, 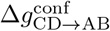, appears in ℋ_conf_:

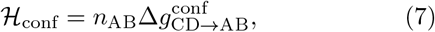

where *n*_AB_ is the number of AB dimers in the assembly. Although the conformational free energy difference 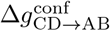 has not been directly estimated from experiments, the results of Mohajerani et al. [35] suggest that 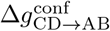 decreases with increasing ionic strength. Consistent with their observations, we vary 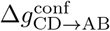 from 5.5*k*_B_*T* to 3.5*k*_B_*T* with increasing ionic strength (see Table I).

The dimer-dimer binding energy function is:

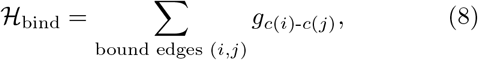

where the binding affinity (free energy) *g*_*c*-*c*′_ depends on the conformations *c, c*′ of the two bound dimers, with values for specific pairs of conformations given in Table I. (To simplify notation, we write e.g. *g*_*c*-*c*′_ = *g*_A-A_ when *c* =BA, *c*′ =AB.) These values were estimated from the buried surface area computed with PDBePISA [84, 85], as reported in Ref. [35]. We report values of *g*_*c*-*c*′_ for each possible interface, relative to a reference value 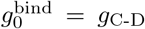 which varies weakly with salt concentration [35], in Table I. Note that the value of 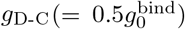 is larger than that used in Mohajerani et al. [35] 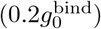. While they found that results were insensitive to the values of *g*_D-C_ in this range, the slightly larger value better reproduces experimental size distributions in the presence of CAMs (Fig. 2) since increasing *g*_D-C_ helps stabilize sheets (indeed, Ref. [35] found that setting 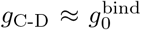 resulted in sheets forming without CAMs). Note also that, unlike in Ref. [35], the values reported in Table I do not account for the entropy of binding; this is instead accounted for via a binding volume, which is built into the KMC algorithm (see SI section S2 B). Together, 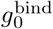 and 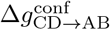 determine the mean dimer-dimer binding affinity within a *T* = 4 capsid [35]:

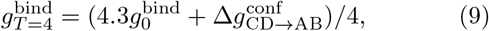

where the factor of 4.3 comes from summing over the energies of the four different contacts present in equal numbers in a *T* = 4 capsid (see Table I). Consistent with experiments, the magnitude of 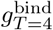 increases with increasing salt concentration (see Table I). We reiterate that, in contrast to Ref. [35], we have not included the binding entropy in these values. We discuss simulation results (Section II) in terms of 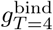 and 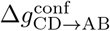.

Finally, the CAM binding energy function is:

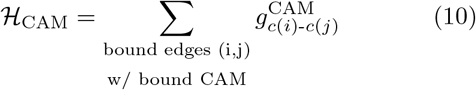

where 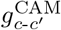 is the binding affinity of CAMs to dimer-dimer interfaces. In our model, CAMs bind only to C-D and D-C interfaces, and hence only 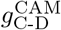 and 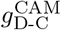 are defined (see Table I). Based on experimental binding affinities of HAP-ALEX (another fluorophore-labeled HAP) to capsids [61], we set 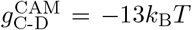. No experimental data exists for binding to D-C interfaces because they do not occur in capsids. We choose 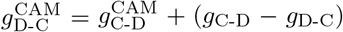, which makes the free energy of a CAM-bound D-C interface equal to that of a CAM-bound C-D interface: 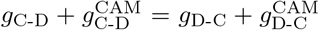 (see SI section S2 A).

### D. Kinetic Monte Carlo (KMC) Move Set

We use a KMC algorithm consisting of nine different types of moves to simulate the assembly dynamics. These moves account for association and dissociation of both (1) single dimers and (2) dimers-of-dimers, (3,4,5) binding and unbinding of dimers within an assembly, which can occur in three topologically distinct ways, (6) dimer conformational switching, binding and unbinding of (7) CAMs and (8) CAM-bound dimers-of-dimers, and (9) thermal fluctuations of the mesh, accomplished via vertex displacement moves (see Figs. S8, S9, and SI Section S2 D for details). KMC moves are proposed with frequencies that reflect the rate constants of each process. A sweep consists of *n*_vert_ attempts to displace a randomly selected vertex, with *n*_vert_ the number of vertices in the assembly, and *n*_dimer_ attempts to change the conformation of a randomly selected edge, where *n*_dimer_ is the number of dimers/edges in the assembly. A sweep sets the timescale of our simulations, *t*_0_ = 10^−5^sec/sweep. We choose this value based on experimental measurements of HBV capsid assembly timescales and previous work [35, 86, 87]. Dimer association/dissociation moves are proposed at a rate 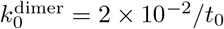, while CAM association/dissociation moves are proposed at a rate 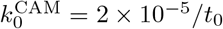. The rate of CAM binding relative to dimer-dimer binding, 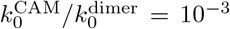, is based on experimental kinetics measurements of HAP-ALEX binding to *T* = 4 HBV capsids [61]. Detailed descriptions of each type of KMC move are given in SI Section S2 D.

### E. Simulation protocol

Each simulation begins with an initial triangular “nucleus” consisting of two AB dimers and one CD dimer as in Ref. [35]. We subsequently run KMC dynamics for a maximum of 2 × 10^8^ sweeps, where we define one sweep as *n*_dimer_ attempted vertex displacement moves. Additionally, simulations end when (i) a complete *T* = 4 or *T* = 3 capsid forms, (ii) assemblies grow to more than 240 edges (twice the size of a *T* = 4 capsid), or (iii) the growth rate of a sufficiently large assembly becomes sufficiently slow (see SI Section S2 C for details). We run 100 independent KMC simulations (using different, randomly-generated, random seeds) for each value of ionic strength, and both with and without CAMs.

### F. Markov state model

To construct a Markov state model (MSM), one chooses a discretization scheme that partitions microscopic configurations into M different ‘microstates,’ such that transitions between microstates are Markovian. Given a collection of many trajectories, one then counts the number of transitions into and out of each microstate after a certain lag time (*τ*) to obtain an *M* × *M* count matrix, ***C***, with elements *C*_*ij*_ representing the number of transitions from microstate *i* to microstate *j* separated by a time *τ*. Column-normalizing the count matrix gives the transition matrix ***T*** (*τ*), with elements given by *T*_*ij*_ = *C*_*ij*_/_*j*_ *C*_*ij*_. The matrix elements *T*_*ij*_ then give the rates of transitioning from microstate *i* to microstate *j*. We partitioned configurations from our KMC trajectories into microstates characterized by the number of AB dimers (*n*_AB_), the number of *T* = 4-like CD dimers 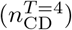, and the number of sheet-like CD dimers 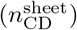. A sheet-like CD dimer forms an edge shared by two triangles (i.e. it is the “central” edge of a diamond), both of which consist solely of CD dimers (see e.g. Fig. 1E). A *T* = 4-like CD dimer forms an edge that is shared by a triangle consisting of only CD dimers and a triangle with two AB dimers and one CD dimer; see Fig. 1Aiii, right.

Given the transition matrix, we can compute the time-dependent probability vector **p**(t) as:

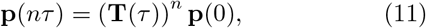

where each element *p*_*i*_(*t*) gives the probability of being in microstate *<u>i</u>* at time *t*, and time is advanced in multiples of the lag time, *t* = *nτ*. We performed a standard test (see SI Section S4 and Fig. S10) to confirm that the MSM dynamics are indeed Markovian for large enough *τ* ; we chose *τ* = 0.02s. As a more stringent test, we also confirmed that the MSM reproduces KMC dynamics: specifically, the MSM yields as a function of time for *T* = 4 capsids and sheets agree well with those obtained from brute-force KMC simulations (Fig. S11).

### G. Committor probabilities

The committor 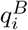 is the conditional probability of reaching a state (or set of states) *B* before reaching another state (or set of states) *A*, given that the system starts in state *i* [52, 53]. For an MSM with transition matrix ***T***, the committor is given by the following equation [52, 53]:

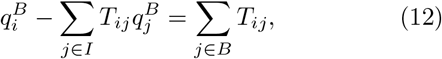

where *I* denotes the set of intermediate states, i.e. those states which are neither in *B* nor in *A*. We compute committors for *T* = 4 assembly 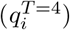 and sheet assembly 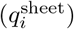. For *T* = 4, state *B* is the microstate with 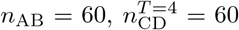, and 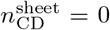. For sheets, state *B* consists of all microstates with 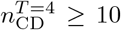. In both cases, state *A* consists of the initial three-dimer nucleus and all absorbing states which are not in state *B*. Eq. 12 is a linear system, which we solve via least squares.

### H. Visualization

Both atomistic structures and coarse-grained simulation snapshots were created using Visual Molecular Dynamics [88].

## ACKNOWLEDGMENTS

We thank Stephen C. Jacobson for providing the experimental data for Fig. 2. This work was supported by the NSF through DMR 2309635 and the Brandeis Center for Bioinspired Soft Materials, an NSF MRSEC (DMR-2011846). JAHP acknowledges funding from NIH P20GM104316-10. Computing resources were provided by the National Energy Research Scientific Computing Center (NERSC), a Department of Energy Office of Science User Facility (award BES-ERCAP0026774); the NSF ACCESS allocation TG-MCB090163; and the Brandeis HPCC which is partially supported by the NSF through DMR-MRSEC 2011846 and OAC-1920147.

## DATA AVAILABILITY

Code for running simulations, as well as code and data for reproducing figures, is available on Zenodo at: [link to be inserted upon publication]

## Appendix A: Simulation parameters

Table I lists the model parameters, their values, and brief descriptions of them.

## Supplementary material

### S1. ADDITIONAL FIGURES

**FIG. S1.**
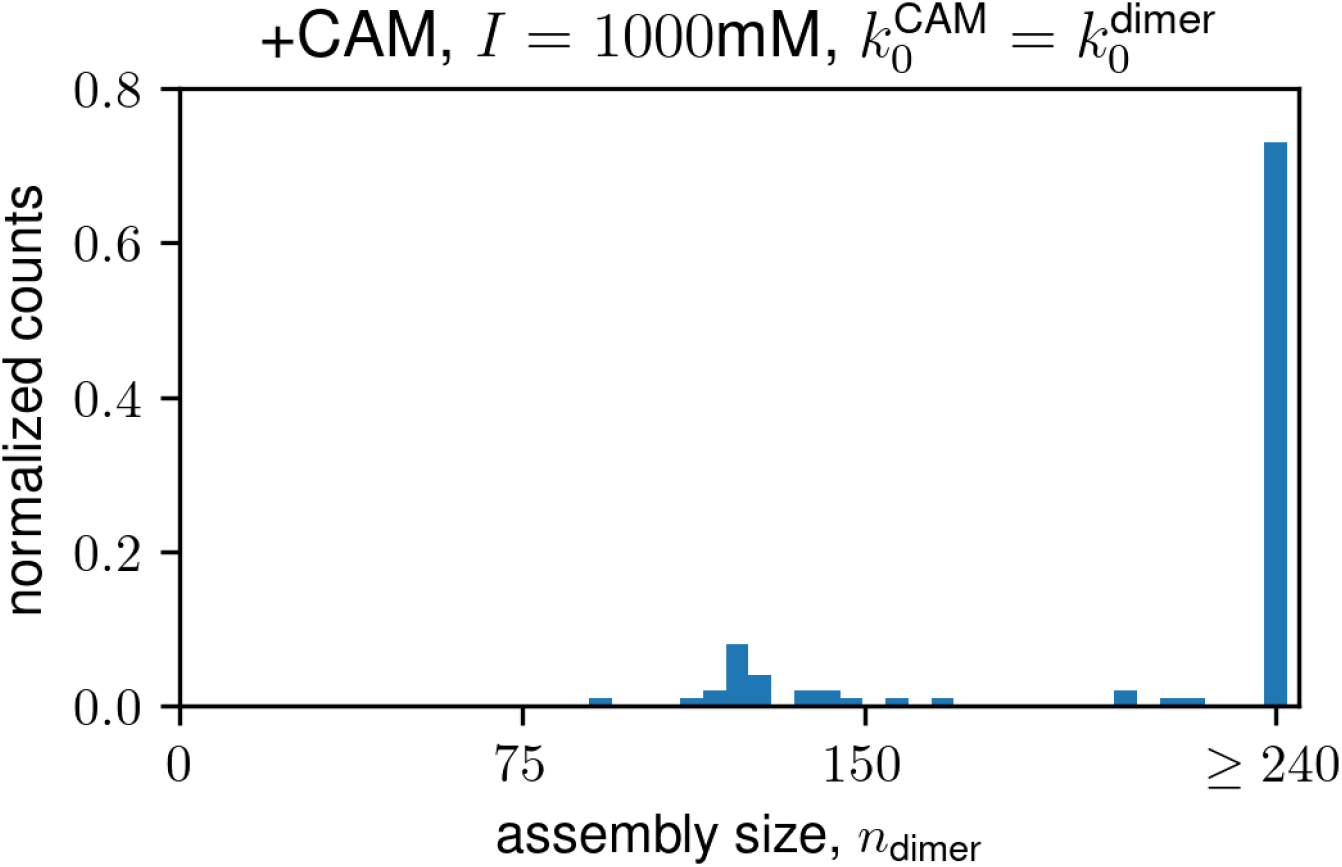
Fast CAM binding yields malformed structures at high salt concentrations. Assembly size distribution from simulations with CAMs at *I* = 1000mM, in which the CAM binding rate constant has been increased from 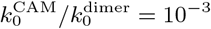 to 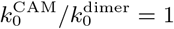. Roughly 70% of the trajectories end in large malformed structures with *n*_dimer_ *≥* 240.

**FIG. S2.**
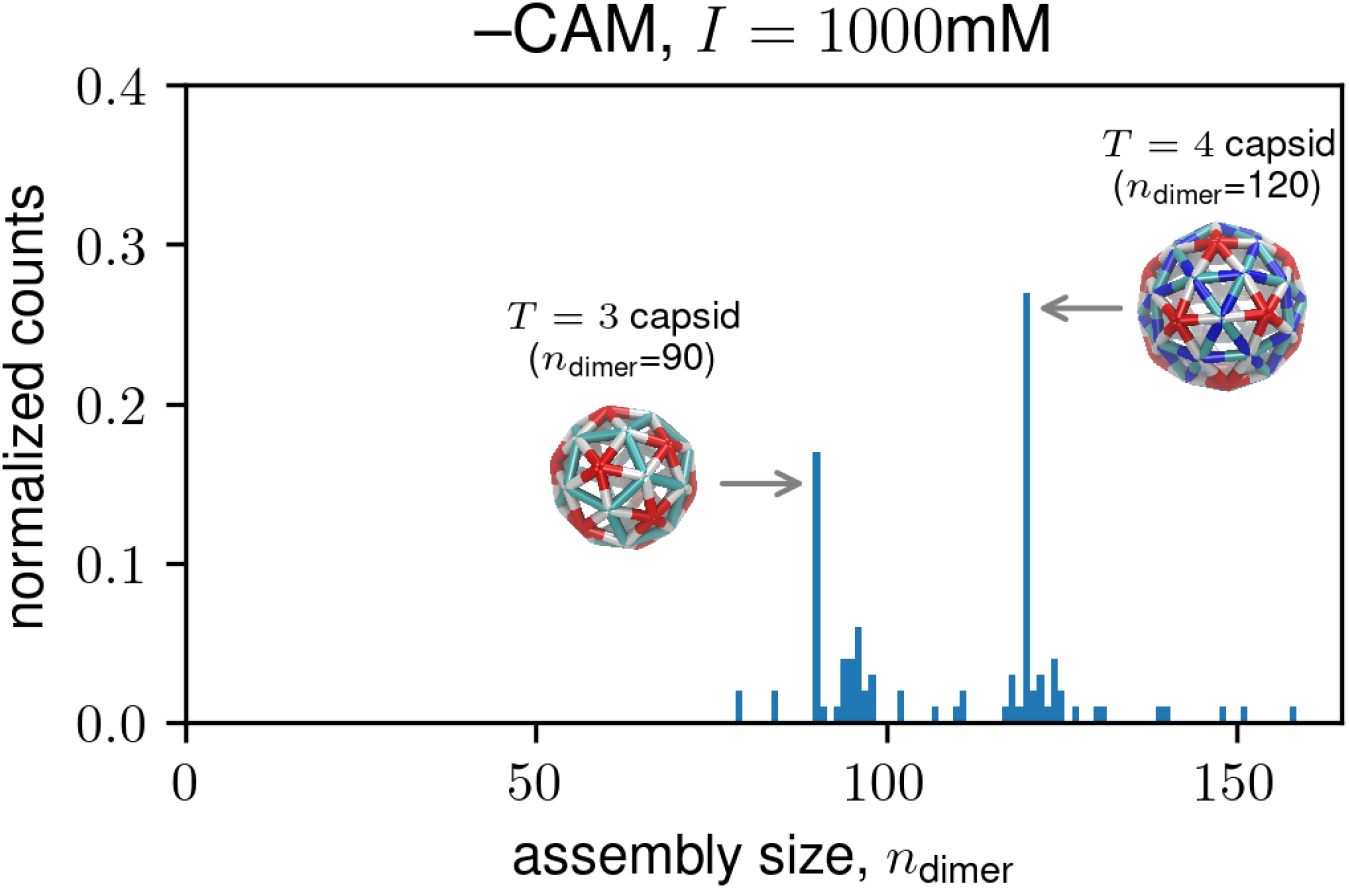
Without CAMs, *T* = 3 capsids occur at high salt concentrations. Assembly size distribution without CAMs at *I* = 1000mM. There are two large peaks at *n*_dimer_ = 90, corresponding to *T* = 3 capsids, and at *n*_dimer_ = 120, corresponding to *T* = 4 capsids. Snapshots show the coarse-grained structures of *T* = 3 and *T* = 4 capsids.

**FIG. S3.**
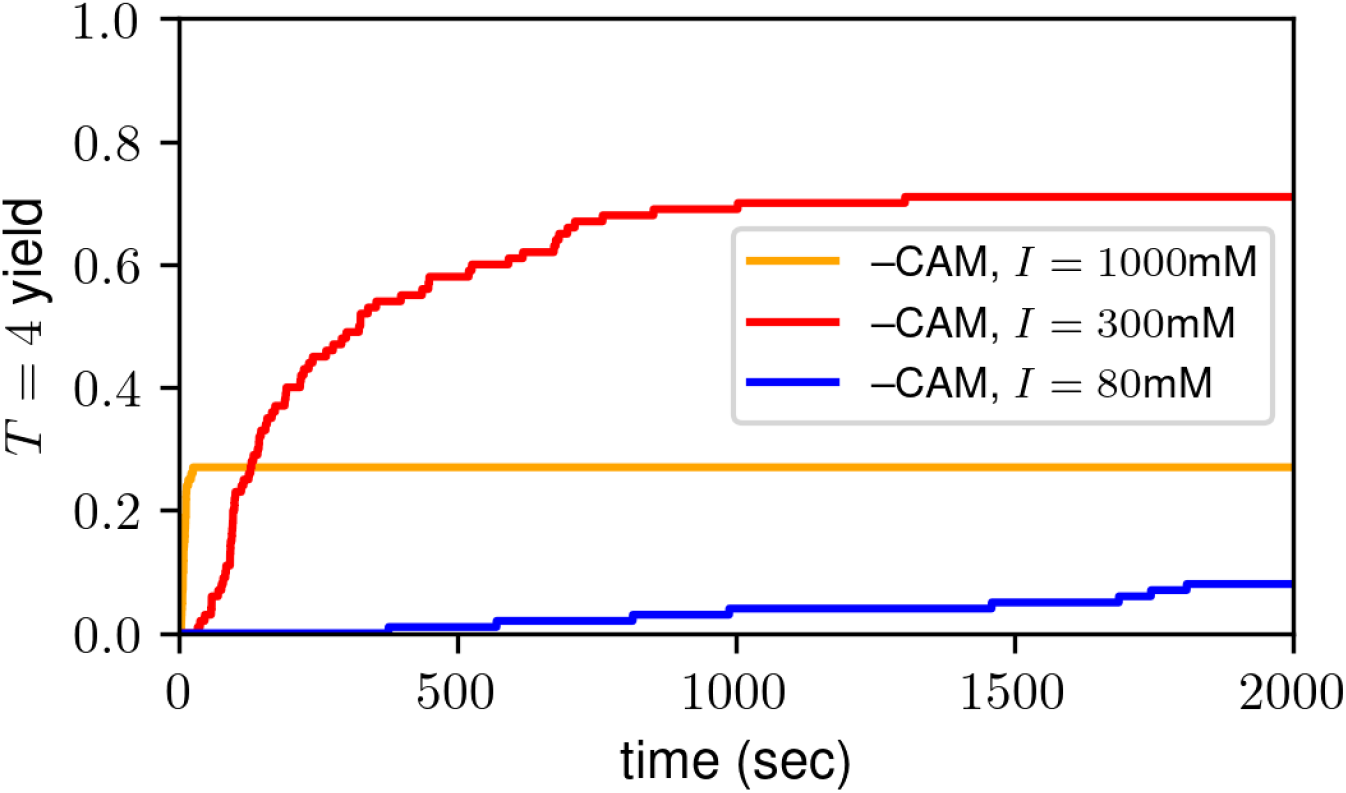
Without CAMs, *T* = 4 capsid formation rate increases with salt concentration. *T* = 4 capsid yield (the fraction of trajectories that end in a *T* = 4 capsid) as a function of time for three different salt concentrations. As the ionic strength increases, the rate at which *T* = 4 capsids form also increases.

**FIG. S4.**
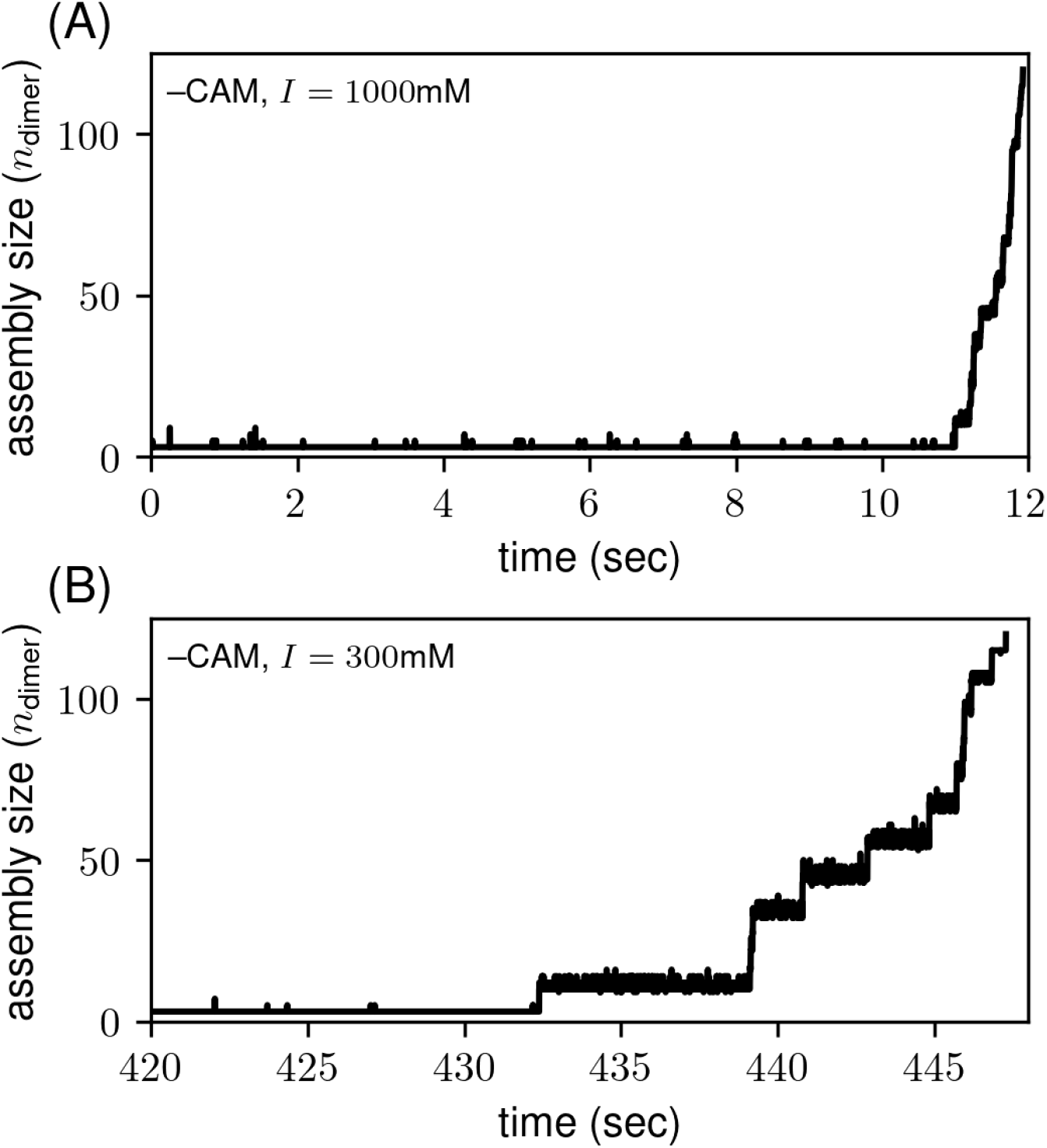
Trajectories leading to *T* = 4 capsids without CAMs are similar to those with CAMs. Assembly size (in *n*_dimer_) versus time for trajectories without CAMs at (A) *I* = 1000mM and (B) *I* = 300mM. Panel A shows the entire trajectory, while the plot in panel B starts at the beginning of the growth phase. These trajectories exhibit similar dynamics and intermediate sizes to the *T* = 4 trajectories with CAMs (Fig. 3 in the main text).

**FIG. S5.**
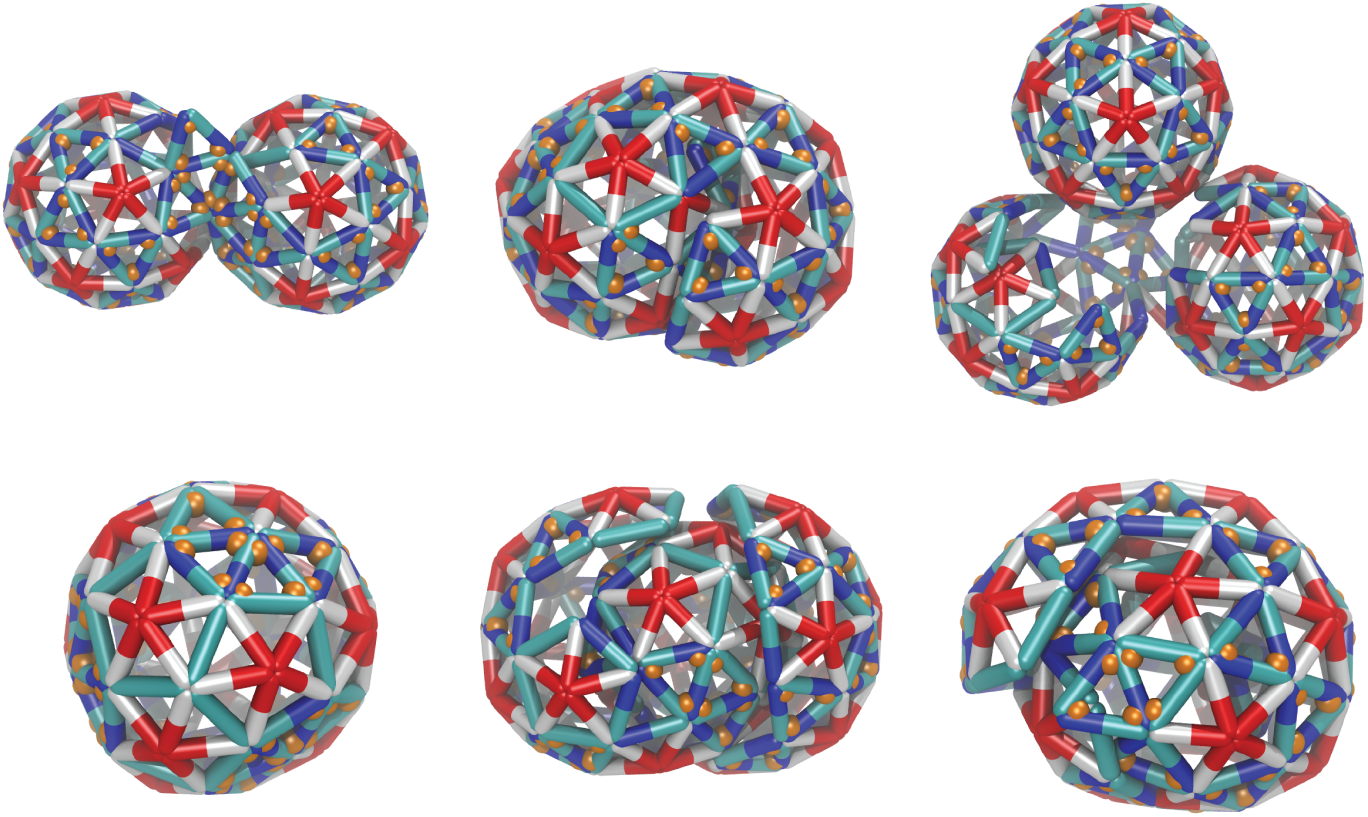
CAMs can induce malformed structures at high salt concentrations. Representative snapshots of malformed structures with CAMs at *I* = 1000mM, including ‘dumbbell’ structures with two connected partial *T* = 4-like shells, ‘triple dumbbell’ structures with three connected partial *T* = 4-like shells, closed shells with non-*T* = 4 symmetry, and shells with overhangs.

**FIG. S6.**
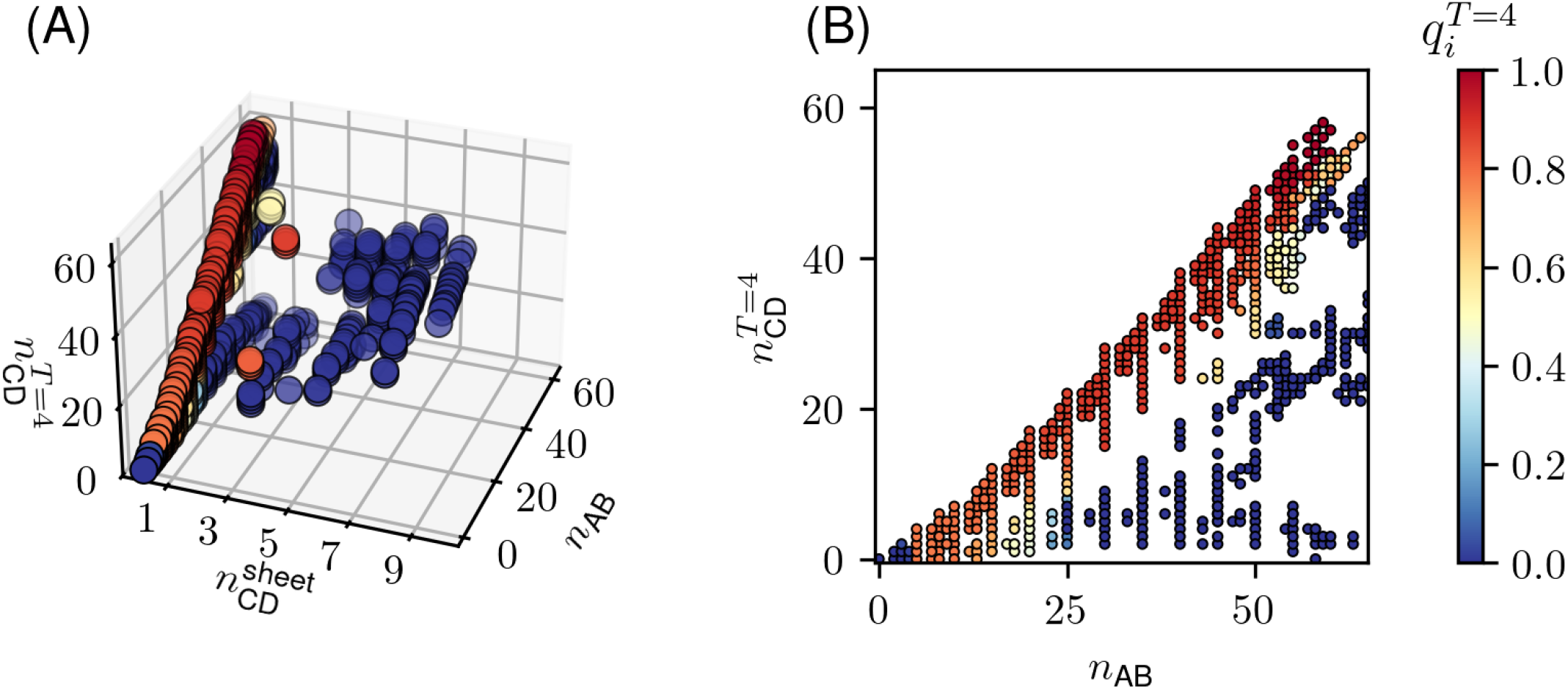
Committors reveal hub states for assembly without CAMs. Committors 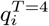 for *T* = 4 capsid assembly (A) as a function of 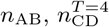, and 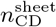 and (B) as a function of *n*_AB_ and 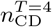 at 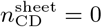. Hub states (with 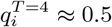) exhibit mixed *T* = 4/*T* = 3 morphologies (see main text Fig. 7D and Ref. [1]).

**FIG. S7.**
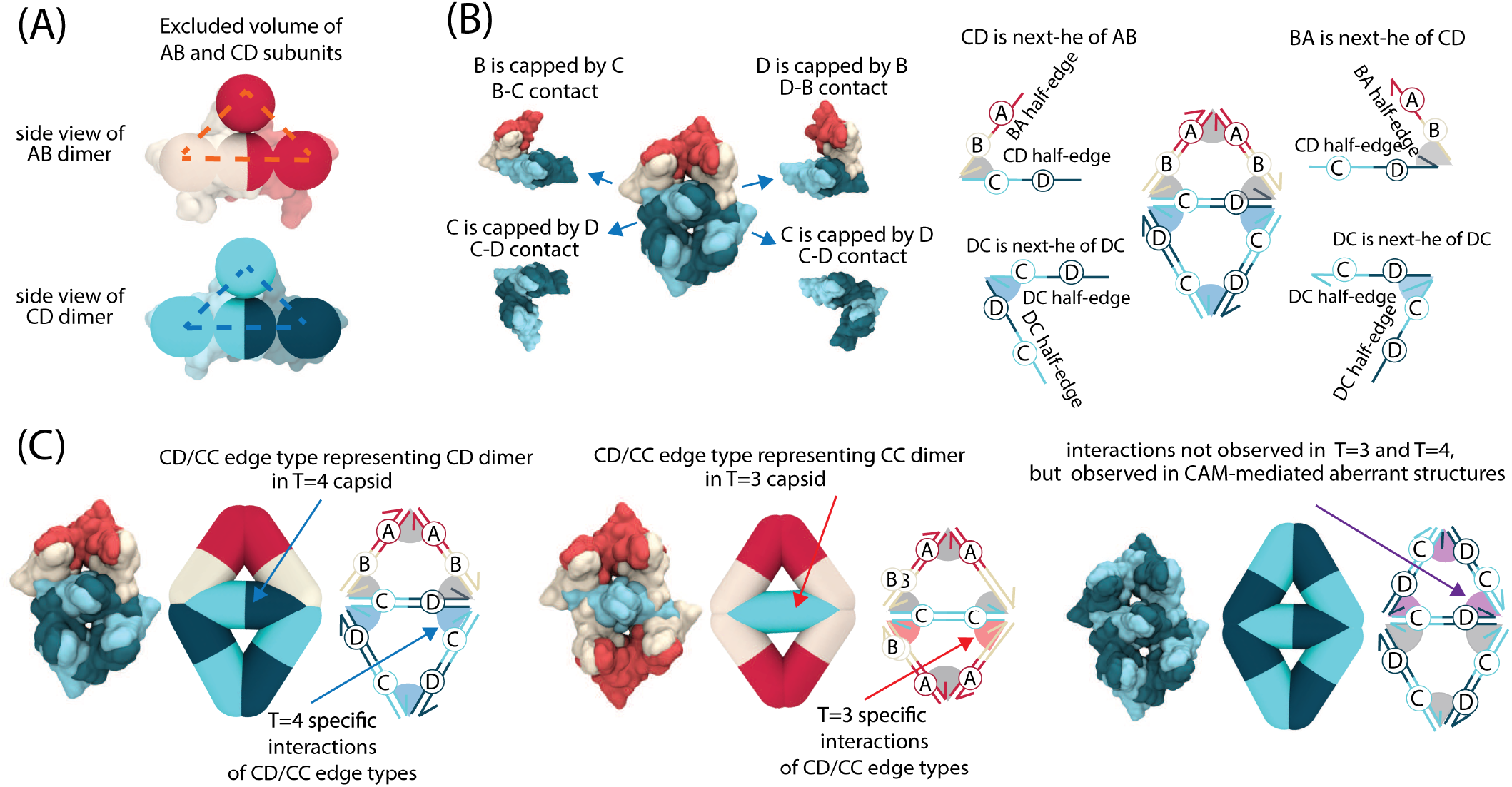
Mapping the half-edge data structure to HBV dimers. **(A)** Side view of overlay of HBV AB and CD dimers and model subunits with excluder pseudoatoms. Subunits are also prevented from overlapping with each other by forbidding the plane shown by dashed lines to intersect the corresponding planes on other subunits. **(B)** The four different contacts of the middle CD subunit in a T=4 intermediate structure (left) and the relevant implementation of each contact by the half-edge data structure (right). In an HBV capsid, each dimer is capped in two of its four contacts, in a specific order as shown in the figure, which is represented by the contacts of two oppositely directed half-edges within an edge. **(C)** The two edge types, AB and CD, are shown along with the different edge interactions that they can make, to represent different conformations and interactions of HBV capsid protein dimers that are observed in available structures: *T* = 4 (left), *T* = 3 (middle), and hexameric sheets (right). Adapted from Ref. [1], copyright 2022 ACS Publications.

### S2. MODEL

Many of the details of our model were described in our previous publication [1]. However, for ease of reference, we have copied those details here, with minor edits. We also describe the implementation of CAMs in the model.

#### A. Implementation of the coarse-grained (CG) model

As described in our previous publication [1], we implement our model of dimer subunits using the half-edge data structure (HE) [2], which is a doubly connected edge list [3]. In our model, each edge corresponds to a protein dimer and consists of two half-edges. There are two edge types: The type AB edge consists of AB and BA half-edges pointing in opposite directions and represents the AB dimer conformation in *T*=3 and *T*=4 capsids; the type CD edge consists of CD and DC half-edges pointing in opposite directions and represents the CD dimer in *T*=4 capsids or the CC dimer in *T*=3 capsids (Fig. S7A,B). Each of these edge types has an equilibrium length 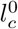 with *c* = AB, CD labeling the conformation, and each interior edge (located between two adjacent triangular faces in the structure) has an equilibrium dihedral angle 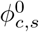 (*s* ∈ (capsid, malformed)). Due to the high structural similarity between CD and CC dimers, we implement them with the same values of 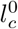 and 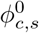 and thus associate them with a single edge type. Throughout the article a CD/CC dimer is called a CD dimer except when it has a (CC/DC)-BA or AB-(CC/DC) interaction (Fig. S7C), since these two interactions are observed in *T*=3 capsids but not in *T*=4 capsids.

The advantage of the half-edge data structure over other, related triangular sheet models [4, 5] is that the two half-edges allow different parameters for protein-protein interactions between different monomer conformations. In particular, with 4 half-edges *c* ∈ {AB, BA, CD, DC}, there are up to 16 different values for binding affinities *g*_*c*-*c*′_ and equilibrium binding angles 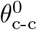 for dimer-dimer interactions. Note that the edges are asymmetric, and thus *g*_*c*-*c*′_ and 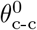 depend on the order of the two half-edges involved; i.e., in general *g*_*c*-*c*′_ ≠ *g*_*c*′-*c*_.

**TABLE S1.** Interaction angles and the relative binding affinities for the different conformations of interacting dimer pairs observed in assembled structures.

| c(h) | c'(next-h) | contact | $g_{c-c'}/g_0^{\text{bind}}$ | $\theta_{c,c'}$ | structure |
| --- | --- | --- | --- | --- | --- |
| BA | AB | A-A | 1.3 | 1.17 | T4,T3 |
| AB | CD(CC) | B-C | 0.9 | 0.98 | T4,T3 |
| CD(CC) | BA | D-B | 1.1 | 0.98 | T4,T3 |
| AB | DC(CC) | B-D | 0.9 | 0.98 | T3 |
| DC(CC) | BA | C-B | 1.1 | 0.98 | T3 |
| DC | DC | C-D | 1 | 1.05 | T4 |
| CD | CD | D-C | 0.5 | 1.05 | w/CAM |

**TABLE S2.** Comparison of dimer-dimer binding affinities in *T*=4 and *T*=3 capsids.

| | $g_{A-A}/k_B T$ | $g_{D-B}/k_B T$ | $g_{B-C}/k_B T$ |
| --- | --- | --- | --- |
| 6UI7 $T=4$ capsid [6] | -8.36 | -7.35 | -6.11 |
| | $g_{A-A}/k_B T$ | $g_{C-B}^{\text{bind}}/k_B T$ | $g_{B-C}/k_B T$ |
| 6UI6 $T=3$ capsid [6] | -8.42 | -7.71 | -6.52 |

Table S1 shows the binding affinity matrix for the possible half-edge-interactions in the model. Each table entry shows the binding affinity (in units of the thermal energy, *k*_B_T) for a pair of half-edges, each belonging to a different edge. The ‘structure’ column lists the structures in which each type of contact is observed. For interactions that involve CC dimers in *T*=3 capsids, CC dimers have similar buried surface area as the CD dimer in *T*=4 capsids in the corresponding local geometry; i.e., CC-BA and AB-CC interfaces in *T*=3 capsids have very similar buried surface areas as CD-BA and AB-DC interfaces in *T*=4 capsids (see Table S2 and reference [6]).

The rest angles 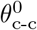 for dimer-dimer interactions observed in *T*=4 and *T*=3 structures were obtained from all-atom (AA) simulations of *T*=4 capsid, as explained in our previous work [1]. All other rest angles are set to *π*/3 since they generally occur in flat or nearly flat hexagonal structures. Similarly, the rest dihedral angles *ϕ*_*c*_ associated with interior edges observed in *T*=4 and *T*=3 structures observed in were obtained from AA simulations. All other equilibrium dihedral angles are set to 0, consistent with flat or nearly flat hexagonal structures.

We represent CAMs as single beads which bind to the interface between two dimers (i.e. where two half-edges in different edges meet). As stated in the main text, we set 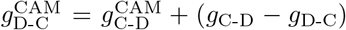, which makes the free energy of a CAM-bound D-C interface equal to that of a CAM-bound C-D interface: 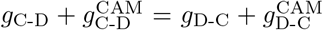 In preliminary simulations where we set 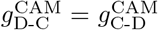, sheets would grow in a slow, step-like fashion, in which CD interfaces formed quickly but DC interfaces formed slowly and transiently (because *g*_C-D_≠ *g*_D-C_). This led to very slow sheet growth on accessible simulation timescales. Setting 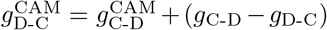 instead resulted in much faster sheet growth and assembly product distributions qualitatively consistent with experiments.

#### B. Monte Carlo (MC) Simulations

Our grand canonical Monte Carlo simulation implementation is adapted from Refs. [1, 5], in which the triangular sheet is represented by edges and vertices. In our model, each edge is associated with two vertices, which together have 2 × 3 degrees of freedom. Any two bound edges in the shell share a vertex, and thus together have 3 × 3 degrees of freedom. For a shell with n_dimer_ edges and n_CAM_ bound CAMs, the grand canonical probability density is

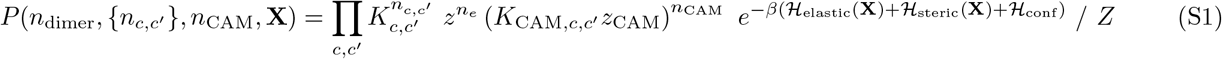

where *c* ∈ {AB,BA,CD,DC} are the different half-edge types (representing their conformations and directions) and n_*c,c*′_ is the number of interactions within the shell configuration between pairs of dimers with *c* and *c*′ conformations respectively. 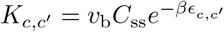, with *ϵ*_*c,c*′_ the binding affinity of interacting half-edges with conformations *c* and *c*′, v_b_ the binding volume, and *C*_ss_ the standard state concentration. *z* = *e*^−*βµ*^ is the activity with the chemical potential *µ* = log(*C*_tot_/*C*_ss_) where *C*_tot_ is the dimer concentration in solution. The standard state concentration is set to *C*_ss_ = 1M. 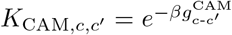, and *z*_CAM_ = *e*^−*βµ*CAM^. The term ℋ_conf_ accounts for the conformational free energy difference of the two types of dimers in the shell. **X** denotes the positions of all the vertices in the sheet, ℋ_elastic_(**X**) accounts for the elastic energy penalty for deviations from the ground state of the sheet, and ℋ_steric_(**X**) accounts for excluded volume among dimers. Finally, the partition function *Z* ensures that the grand canonical probability density is normalized.

The binding volume 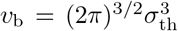, where 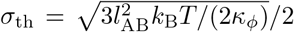 is a thermal length scale. The quantity log(v_b_C_ss_) represents an entropic free energy penalty for dimer-dimer binding, effectively modifying the free energy *g*_*c*-*c*′_ relative to its “bare” value *ϵ*_*c,c*′_, i.e. *<u>g</u>*_*c*-*c*′_ = *ϵ*_*c,c*′_ − *k*_B_T log(*v*_b_*C*_ss_).

In each MC step, a trial Monte-Carlo move ν is chosen randomly from the list of Monte-Carlo moves (described below) according to its relative trial rate 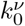. The trial moves are accepted/rejected based on the Metropolis-Hastings algorithm, ensuring detailed balance.

#### C. Shell assembly simulations

To simulate capsid assembly dynamics, we intersperse a variety of MC moves, described in the next section, that are designed to capture the physical dynamics of assembling subunits. Each simulation begins with an initial state comprising three edges in a triangular face. This state is known to be a highly populated intermediate, and is estimated to be the critical nucleus for HBV capsid assembly [7–9]. Each simulation contains a single assembly, which undergoes exchange of subunits and CAMs with a reservoir according to the grand canonical probability density, Eq. (S1). The simulations are performed for a maximum of 2 × 10^8^ sweeps, where a sweep is defined as a set of trial moves consisting of addition/removal, binding/unbinding, shell relaxation, and conformational switch moves such that each edge on average will have undergone one conformational switch move and each vertex on average will have undergone one vertex displacement move. Simulations are stopped early if the assembly forms a closed structure, defined as a structure in which every edge has its maximum number of four interactions. Simulations are also stopped early if the assembly becomes stalled in a sufficiently long-lived intermediate, defined as a structure with *n*_dimer_ ≥ 75 edges which has grown by no more than two dimers in the previous *n*_stall_ = 100/α sweeps, where α is the average growth rate (difference between the number of dimers every 10^4^ MC sweeps, averaged over subsequent intervals of 10^4^ MC sweeps) in a given simulation after the structure has reached *n*_dimer_ = 35 edges.

#### D. MC moves

The set of MC moves and their acceptance criteria are described here. Moves are accepted or rejected according to the Metropolis-Hastings acceptance criteria [10, 11]:

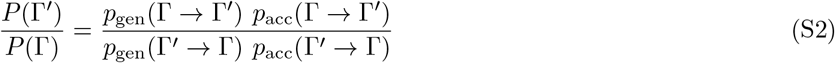

where Γ and Γ′ denote the initial and trial states, the probability of a state with *n*_dimer_ edges and *n*_CAM_ CAMs is 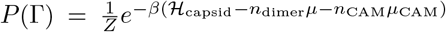 with *Z* the grand canonical partition function, and *p*_gen_ and *p*_acc_ are respectively the probabilities for generating and accepting trial moves.

##### Notation

Figure S8A shows a schematic of an example structure, with explanations of some notation that will be used in the following descriptions of the MC moves. Edges which have their full complement of four interactions are denoted as *non-boundary edges*. The two half-edges within a non-boundary edge are denoted as *non-boundary half-edges* (black arrows in Figure S8A). Each non-boundary half-edge interacts with a half-edge on each of the two neighboring edges, which are denoted as *next-h* (from its head) and *prev-h* (from its tail). An example of a half-edge **h** with its next-h and prev-h is shown in Figure S8A.

A *boundary half-edge* has at least one unbound end; i.e., it lacks an interaction with at least one of its adjoining half-edges (next-h or prev-h). By keeping track of the set of boundary half-edges, the simulation algorithm is able to efficiently choose possible trial moves which involve binding new edges. In the MC implementation used for this work, we allow only one half-edge in each edge to be a boundary half-edge, which prevents formation of dangling edges (that have only one interaction) and star-like configurations. Any trial move that results in formation of an edge that comprises two boundary half-edges is rejected.

In the description of the MC moves that follows we describe moves in terms of changes in half-edges. However, note that in general each move affects both half-edges within an edge, and these effects occur simultaneously.

##### Vertex move

A vertex is randomly chosen, and a trial <u>is made to displ</u>ace it by the vector 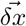, whose components are chosen from a Gaussian distribution 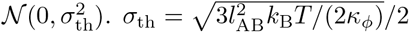 is the length scale for the thermal fluctuations of the system. In this move, the numbers of dimers, CAMs, dimer-dimer and dimer-dimer-CAM interactions, vertices and edge types are unchanged (Fig S8B). The move is accepted with probability:

**FIG. S8.**
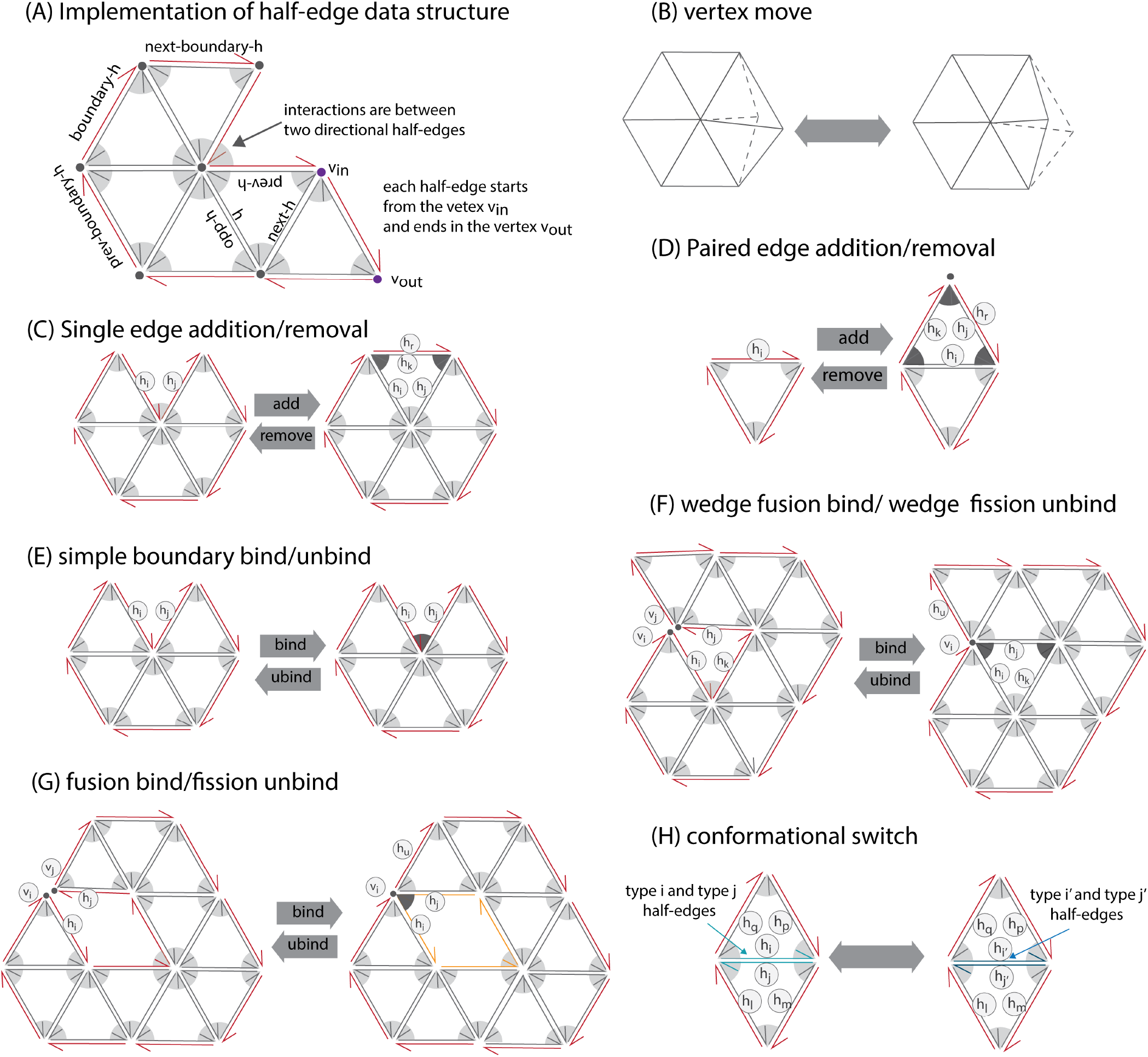
Schematic of the half-edge implementation and the MC moves. **(A)** Half-edge implementation. Each edge in the model is composed of two half-edges. Half-edges with no open ends are designated as *non-boundary half-edges*, and drawn as grey arrows. For a compact notation, for a given half-edge, the half-edges that it interacts with on its tail and on its head are designated as *previous-h* and *next-h* respectively. Half-edges that are open (have no interaction partner) on their head and/or tail are designated as *boundary half-edges*, and drawn as red arrows. Each boundary half-edge has a *next-boundary-h* and a *prev-boundary-h*, but there is not necessarily an interaction between a half-edge and its next-boundary-h or prev-boundary-h. **(B)** The *vertex move*, in which a vertex is randomly displaced. **(C)** *Single edge addition/removal*. The single edge addition move adds an edge (two half-edges) to the boundary of the shell, if the two selected boundary half-edges are bound to each other. This results in two new interactions, drawn as dark gray wedges in the schematic. The reverse move, a single edge removal, breaks two interactions and removes two half-edges form the shell. **(D)** The *paired edge addition/removal* move adds/removes two edges to the shell, resulting in forming/breaking three interactions (dark gray wedges in the schematic). **(E)** The *simple boundary bind/unbind* makes/breaks an interaction between two boundary half-edges. **(F)** *Wedge fusion bind / wedge fission unbind*. A wedge fusion move adds an interaction between two boundary half edges that are sufficiently near each other. In the example shown, in the left configuration there are two nearby boundary half-edges that can bind. This causes a vertex to be removed and adds two new interactions. The reverse move, wedge fission unbinding, results in adding a new vertex and breaking two interactions. **(G)** *Fusion bind / fission unbind*. In fusion binding, two close boundary half-edges are bound, resulting in removal of a vertex and one new interaction (dark grey wedge in the schematic). The reverse move, fission unbinding, results in removal of one vertex and breakage of one interaction. **(H)** The *conformational switch* move changes the conformations of the two half-edges within one edge. Adapted from Ref. [1], copyright 2022 ACS Publications.

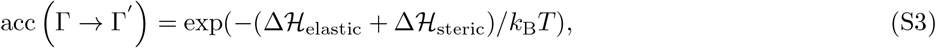

and the generation probability is gen (Γ → Γ′)= 1.

##### Single edge addition/removal

A half-edge **h**_i_ is randomly chosen from the boundary half-edges. If **h**_i_ is bound to another half-edge **h**_j_ from one end, a trial is made to add a new edge, consisting of two half-edges, **h**_k_ and its opposite half-edge, connecting the other end of **h**_i_ to **h**_j_ (Fig S8C). The generation probability is

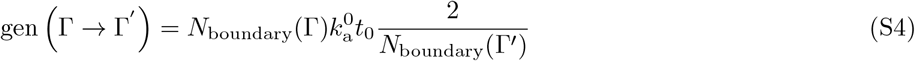

where *N*_boundary_ is the number of boundary half-edges, 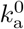 is the trial-move rate for addition/removal moves, and *t*_0_ is the simulation timescale, as explained in the main text.

The newly added half-edge **h**_k_ interacts with **h**_i_ and **h**_j_, with conformation-dependent binding affinities, so the grand canonical probability of the new configuration relative to Γ is

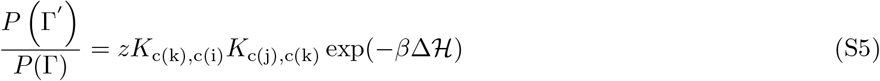

where c(i), c(j), and c(k) are the conformations of half-edges **h**_i_, **h**_j_, and **h**_k_ respectively and Δ ℋ = Δ ℋ_elastic_ + Δ ℋ_steric_ + Δ ℋ_conf_ involves volume exclusion, the elastic energy of the new edge and its bound edges, and the conformational free energy of the new edge.

For removal of a single edge, a half-edge **h**_r_ is randomly chosen from the boundary half-edges. A trial is made to remove **h**_r_ and its opposite half-edge **h**_k_, which includes unbinding **h**_k_ from its next-h **h**_i_ and prev-h **h**_j_. The generation probability of edge removal is:

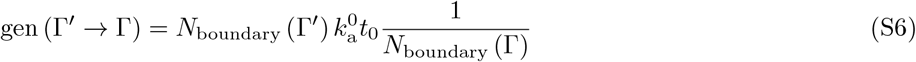

The acceptance criterion is:

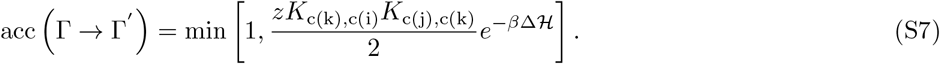

##### Paired edge addition/removal

In our model, edge additions that result in configurations with a dangling edge (which has only one interaction) are followed by addition of a second edge that closes the triangle. This choice is made because, under productive assembly conditions, dangling edges are highly unstable and quickly dissociate. Thus, simulations would spend the majority of their time on consecutive additions and removals of dangling edges. In previous Brownian dynamics simulations [12] we observed that in such situations net growth of assemblies was usually associated with either association of oligomers, or the rapid succession of additions of more than one subunit [12]. A similar conclusion was made from kinetic MC simulations [13]. To allow for this possibility, we include a move that enables additions and removals of dimers-of-dimers.

The two consecutive edge additions are attempted as follows: A half-edge **h**_i_ is randomly chosen from the boundary half-edges. If it is open on both ends (i.e. it has neither next-h nor prev-h), a trial is made to add two new edges that bind to the selected edge. This move also includes adding a new vertex to the shell, at the intersection of the two newly added edges. To specify the coordinate of the new vertex, first the equilibrium position of the new vertex 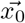 is selected based on the conformations of **h**_i_, **h**_j_ and **h**_k_. The new vertex is then displaced to 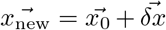 where the components of 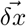 are selected from the Gaussian distribution 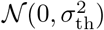 with *σ*_th_ described above. The generation probability is

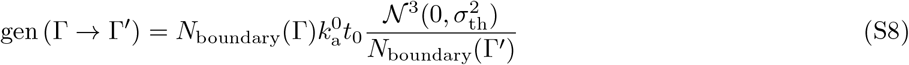

where 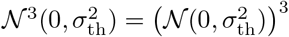.

The first added half-edge (**h**_j_) makes a new interaction with one end of **h**_i_, and the second added half-edge (**h**_k_) interacts with the open ends of **h**_j_ and **h**_i_.

For the reverse move, a half-edge **h**_r_ is randomly chosen from the boundary half-edges. If removal of this half-edge (and its opposite half-edge) results in a dangling edge, the dangling edge will also be removed. In this move, two edges, three interactions, and a vertex are removed. The generation probability for removal is:

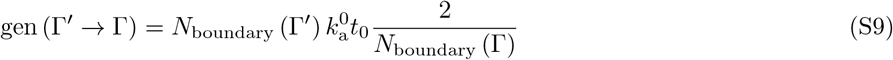

and the acceptance criterion for addition is:

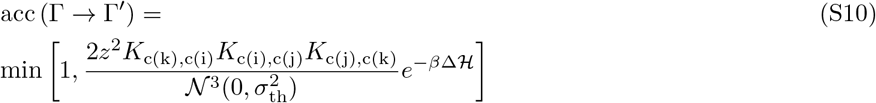

where again Δℋ = Δℋ_elastic_ + Δℋ_steric_ + Δℋ_conf_ involves volume exclusion, the elastic energy of the new edges and their bound edges, and the conformational free energy of the new edges.

##### Simple boundary binding/unbinding

In a simple boundary binding, we attempt to make a new interaction between two edges whose ends are nearby but unbound to each other (Fig. S8E). For this move, a half-edge **h**_i_ is randomly chosen from the boundary half-edges. If **h**_i_ is open on both ends (has neither next-h nor prev-h) and makes a wedge with the next-boundary-h or prev-boundary-h **h**_j_, (i.e., the angle between the two edges θ < π/2) an attempt is made to bind **h**_i_ to **h**_j_.

For the opposite process, simple boundary unbinding, an edge is chosen randomly from the boundary edges and if it has a next-h or prev-h (it cannot have both since it is a boundary edge), an attempt is made to unbind. The acceptance criterion for simple boundary binding is:

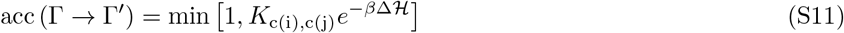

Here Δℋ only involves the excluded volume and elastic energies.

##### Wedge fusion binding/Wedge fission unbinding

For a wedge fusion move, a half-edge **h**_i_ is randomly chosen from the boundary half-edges. A wedge fusion is attempted if: 1) **h**_i_ makes a wedge with angle α < π/2 with another boundary half-edge **h**_j_, 2) v_in_ of **h**_j_ is within δx_fuse_ of v_out_ of **h**_i_, 3) the next-boundary-h of **h**_i_ (**h**_k_ in Fig S8F) is the same as prev-boundary-h of **h**_j_, and 4) **h**_k_ is bound to **h**_j_ or **h**_i_. For the implementation of this attempt, **v**_i_ and **v**_j_ and their associated edges are fused to a new vertex **v**_k_ at the midpoint between **v**_i_ and **v**_j_. Two new interactions are made: **h**_i_ is bound to **h**_j_ and the third half-edge in the triangle **h**_k_ is bound to **h**_i_ (or to **h**_j_).

For a wedge fission unbinding, a vertex **v**_k_ is chosen at one end of a randomly chosen boundary half-edge. An interaction of that vertex between an associated half-edge **h**_i_ and its next-h **h**_j_ is randomly chosen for the wedge fission unbinding attempt. Then, v_out_ of the incoming half-edge **h**_i_, with all its associated edges, is moved to the new vertex **v**_i_ at 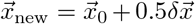; and v_in_ of the outgoing half-edge **h**_j_, with all its associated edges, is moved to the new vertex **v**_j_ at 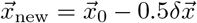, where the components of 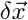 are chosen from the Gaussian distribution 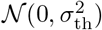. To maintain the proper topology of the shell, the interaction of the third half-edge in the triangle (randomly chosen as **h**_k_-**h**_i_ or **h**_k_-**h**_j_) is also removed. The acceptance criterion for wedge fusion binding is:

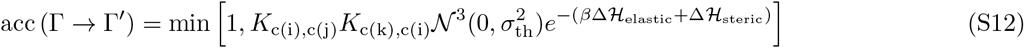

##### Fusion binding / fission unbinding

For a fusion binding move, a half-edge **h**_i_ is randomly chosen from the boundary half-edges. The move is attempted if: 1) **h**_i_ forms a wedge with angle *α* < *π*/2 with another boundary half-edge **h**_j_, 2) v_in_ of **h**_j_ is within δx_fuse_ of v_out_ of **h**_i_, and 3) the next-boundary-h of **h**_i_ (**h**_k_ in Fig S8G) is not the same as prev-boundary of **h**_j_. Similar to the wedge binding move, two vertices and their associated edges are fused into a new vertex **v**_k_ at the midpoint between **v**_i_ and **v**_j_. This move results in an additional boundary loop in the structure (orange loop in Fig S8(G) right) and the bound vertex will be a double-boundary vertex, meaning that it is shared between two boundary loops. A boundary loop can be found by starting from a random boundary half-edge, and moving along the next-boundary-h elements until returning to the original half-edge.

A fission unbinding move is attempted if there is more than one boundary loop in the structure. A vertex **v**_k_ is chosen at one end of a randomly chosen boundary half-edge. If this vertex is a double-boundary vertex, the fission unbinding move will be attempted, by splitting the edges ending in **v**_k_, to form to vertices **v**_i_ and **v**_j_ (similar to the wedge fission unbinding move), and merging the two boundary loops. The implementation of adding the new vertex to the shell is similar to the wedge fission unbinding move, except that there is only one unbinding event (**h**_i_-**h**_j_).

The acceptance criterion for fusion binding is:

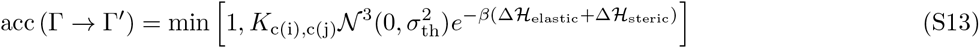

##### Conformational switch

A half-edge **h**_i_ is randomly chosen from the set of all edges in the structure, and a trial is made to change the conformation of **h**_i_ from c(i) to *c*′(i) and the conformation of its opposite half-edge **h**_j_ from c(j) to *c*′(j). The generation probability is gen (Γ → Γ′) = 1, and the move is accepted with probability:

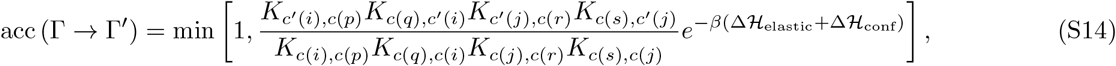

where p and q are the indices of the next-h and prev-h of i, and r and s are the indices of the next-h and prev-h of j.

##### CAM binding/unbinding

For a CAM binding move, a half-edge **h**_i_ is randomly chosen. If the half-edge is type CD or DC and has a prev-h, and no CAM is bound already, then an attempt is made to add a CAM to the interface between **h**_i_ and prev-h (for the purposes of bookkeeping, we associate the CAM with **h**_i_ rather than prev-h; see Fig. S9A). The generation probability is:

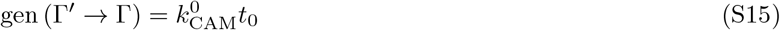

The move is accepted with probability:

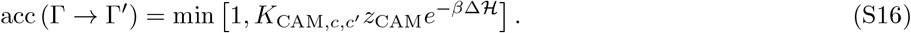

Similarly, for a CAM unbinding move, a half-edge is randomly chosen. If the half-edge is type CD or DC and a CAM is bound to its interface with prev-h (if prev-h exists), then an attempt is made to remove the CAM from the interface. The generation probability for such moves is:

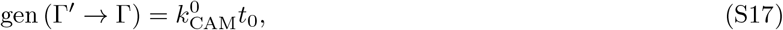

and the acceptance probability is:

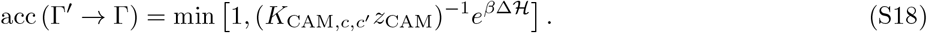

In principle, the procedure detailed above could result in CAMs being bound to the interface between CD/DC and AB/BA dimers. However, because we set *g*^CAM^ = 0 for any interfaces that are not C-D or D-C, in practice CAM is only ever observed bound to C-D and D-C interfaces.

##### CAM-bound dimer-of-dimers addition/removal

We also allow for the binding and unbinding of CAM-bound CD dimers-of-dimers (Fig. S9B). The rules are the same as for paired edge addition/removal; one simply accounts for the additional energy of the CAM bound to the dimer-of-dimers interface. Importantly, dimers-of-dimers cannot be removed if CAMs are bound to either of the other dimer-of-dimers interfaces in the triangle to which the selected half-edge belongs (see Fig. S9C).

**FIG. S9.**
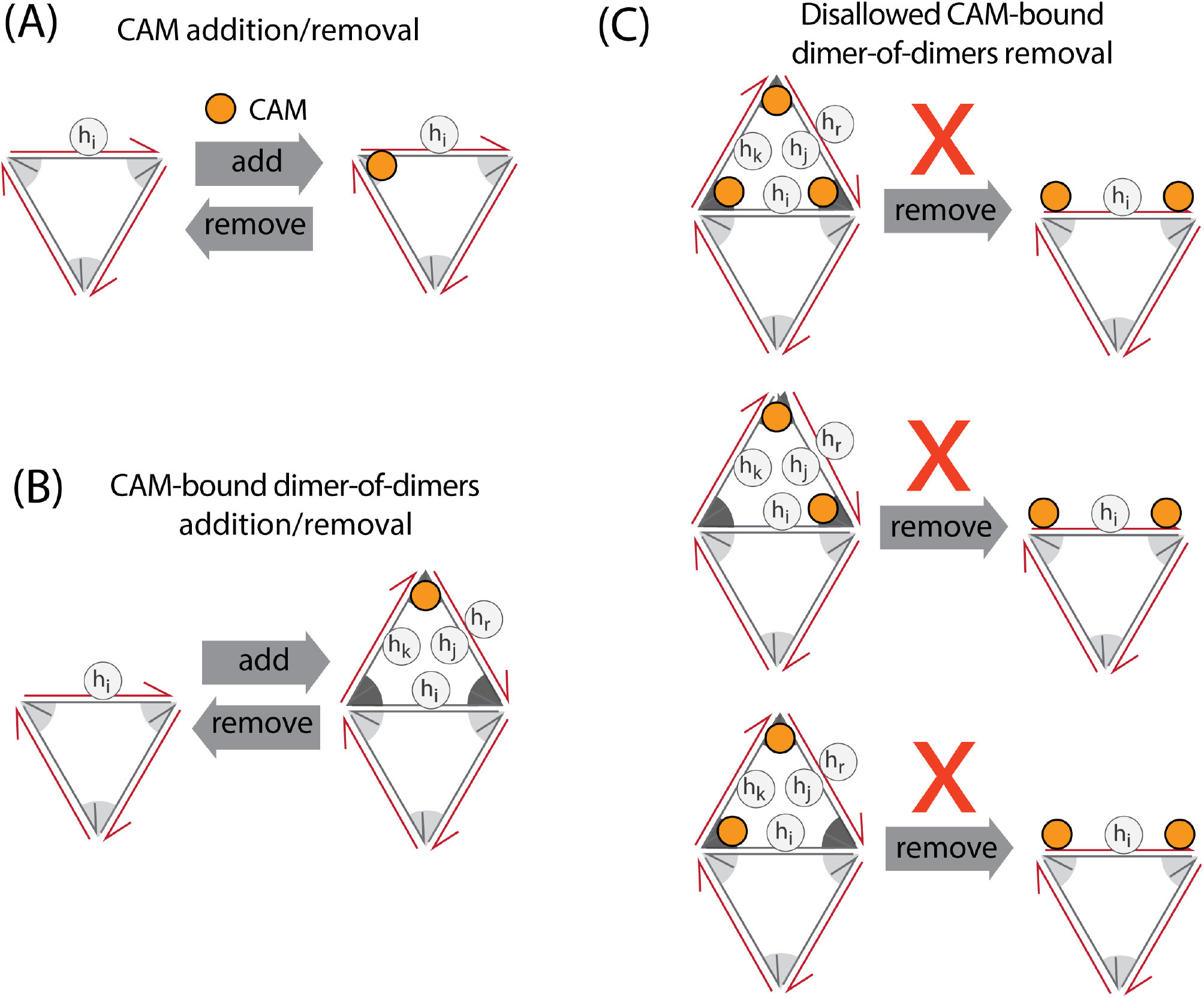
Schematic of MC moves involving CAMs. **(A)** The *CAM addition/removal* move adds/removes a CAM to/from the interface between two bound edges. **(B)** The *CAM-bound dimer-of-dimers addition/removal* move adds/removes two edges with a CAM bound to their interface to the shell, resulting in forming/breaking three interactions (dark gray wedges in the schematic). **(C)** CAM-bound dimer-of-dimers removal moves that would result in CAMs on boundary half-edges are disallowed.

**TABLE S3.**
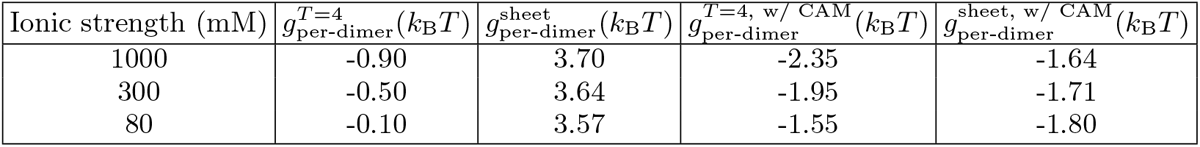
Values of per-subunit binding free energies, with and without CAMs, at different ionic strengths.

### S3. FREE ENERGIES OF CAPSIDS AND SHEETS

The per-dimer free energy of *T*= 4 capsids and sheets depends upon both ionic strength and the presence of CAMs.

Without CAMs, the per-dimer free energies of *T*= 4 capsids 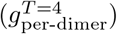 and sheets 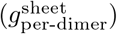 are given by:

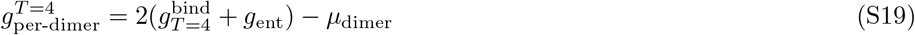

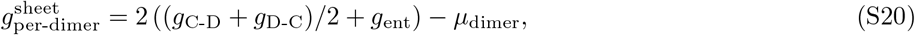

where *g*_ent_ ≈ 3.42*k*_B_T [1] is the entropic penalty for dimer-dimer binding and the factors of two account for the fact that each dimer in a capsid is bound to four other dimers. The expression for 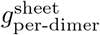 neglects the cost associated with the boundary of the sheet. Alternatively, if CAMs are bound to each possible site in the structures, the per-subunit free energies are:

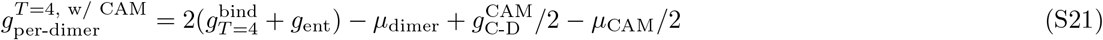

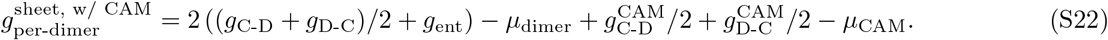

Values of 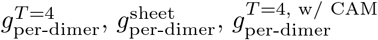, and 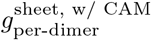 for different ionic strengths are shown in Table S3.

**FIG. S10.**
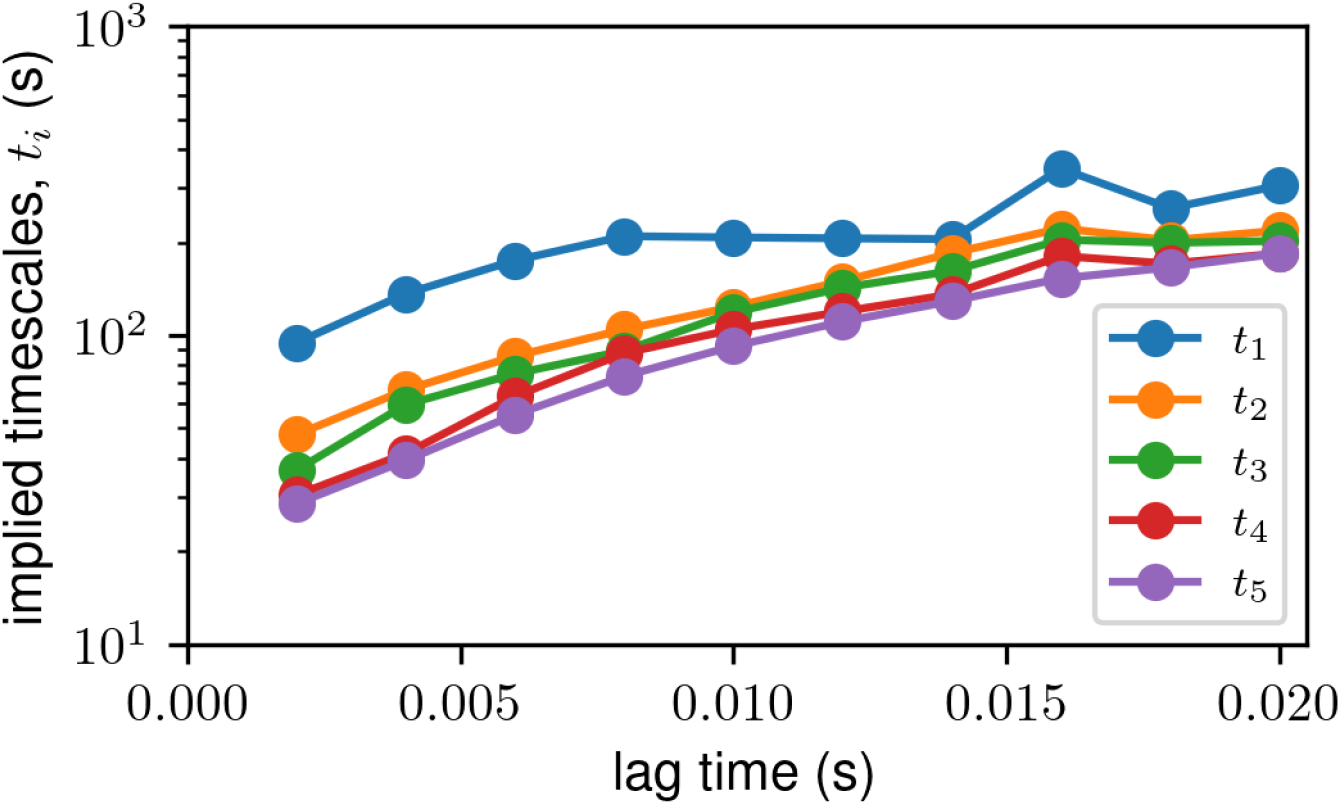
Implied timescales vary slowly with lag time. Plot showing implied timescales *t*_*i*_ = − *τ/* log *λ*_*i*_ computed by diagonalizing the MSM transition matrix versus the lag time (*τ*) in seconds. We show the five longest timescales in different colors, corresponding to the five highest eigenvalues *λ*_*i*_ of the transition matrix with *λ*_*i*_ *<* 1.

### S4. MARKOV STATE MODEL VALIDATION

To validate our Markov state model (MSM), we perform two tests. In the first test (widely used in the MSM literature, e.g. [14]), we compute implied timescales of the MSM for varying lag times *τ*. Implied timescales t_*i*_ are defined in terms of the eigenvalues λ_*i*_ of the MSM transition matrix ***T*** (*τ*):

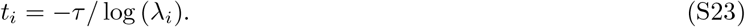

If the dynamics are Markovian, then *t*_*i*_ should be independent of *τ*. In practice this is almost never exactly true, and instead a rule of thumb is that *t*_*i*_ should vary only slowly with *τ* for the largest implied timescales (i.e. largest eigenvalues with λ_*i*_ < 1). Fig. S10 shows that this is indeed the case for the largest five implied timescales in our MSM.

As a second test, we compute the yields of *T*= 4 capsids and sheets as a function of time with the MSM, and compare them to yields computed directly from KMC simulations. This is a much more stringent test of the MSM than the implied timescale test, because it requires that all elements of the transition matrix be accurately estimated. The MSM yield versus time is given by *∑*_*i*∈*B*_ *p*_*i*_(*t*), where the sum is over microstates i associated with the product state B (*T*= 4 capsid or sheet) and p_*i*_(t) is computed via Eq. 11 in the main text. The KMC yield is the fraction of trajectories in which a product (*T*= 4 capsid or sheet) has formed. Fig. S11 shows that the MSM yield dynamics agrees very well with that of the brute-force KMC simulations.

**FIG. S11.**
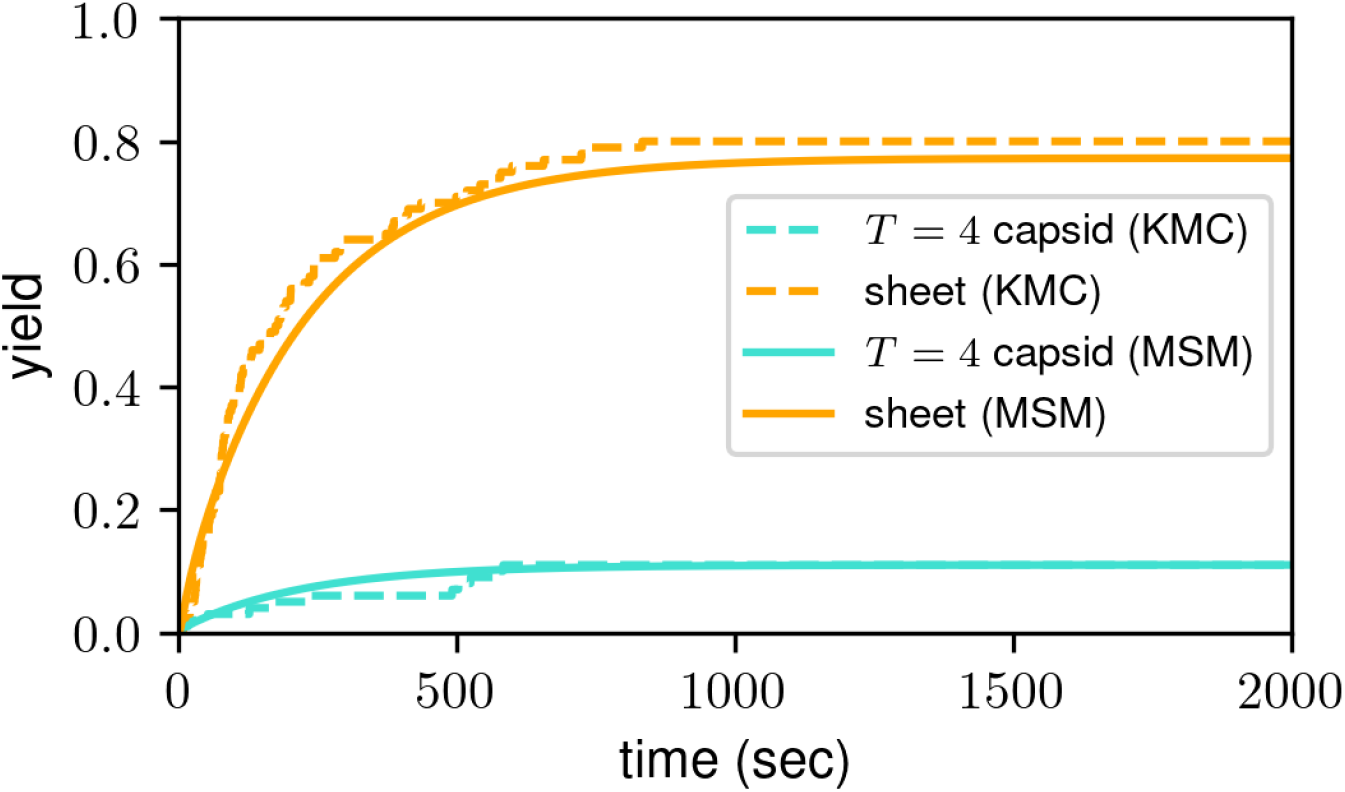
MSM dynamics agree well with KMC dynamics. Yield, i.e. the fraction of trajectories ending in either *T*= 4 capsids (turquoise) or sheets (orange), versus time. The dashed lines show results from KMC simulations, while the solid lines show results from the MSM (obtained by solving Eq. 11 in the main text).

### S5. MOVIE DESCRIPTIONS

- **Movie S1**: Trajectory showing *T*= 4 capsid assembly with CAMs at *I* = 1000mM. The corresponding assembly size versus time plot is shown in the main text (Fig. 3A).
- **Movie S2**: Trajectory showing malformed assembly with mixed *T*= 3/T = 4 morphology with CAMs at *I* = 1000mM. The corresponding assembly size versus time plot is shown in the main text (Fig. 3B).
- **Movie S3**: Trajectory showing *T*= 4 capsid assembly with CAMs at *I* = 300mM. The corresponding assembly size versus time plot is shown in main text (Fig. 3C).
- **Movie S4**: Trajectory showing malformed assembly with mixed sheet/T = 4 morphology with CAMs at *I* = 300mM. The corresponding assembly size versus time plot is shown in the main text (Fig. 3D).
- **Movie S5**: Trajectory showing sheet assembly with CAMs at *I* = 80mM. The corresponding assembly size versus time plot is shown in the main text (Fig. 3F).
- **Movie S6**: Trajectory showing *T*= 4 capsid assembly without CAMs at *I* = 300mM.

## Notes

### Competing Interest Statement

The authors have declared no competing interest.

## References

[1] M. Krupovic and E. V. Koonin, Multiple origins of viral capsid proteins from cellular ancestors, Proceedings of the National Academy of Sciences 114, E2401 (2017).

[2] D. M. Salunke, D. L. Caspar, and R. L. Garcea, Poly-morphism in the assembly of polyomavirus capsid protein VP1, Biophysical Journal 56, 887 (1989).

[3] L. S. Ehrlich, B. E. Agresta, and C. A. Carter, Assembly of recombinant human immunodeficiency virus type 1 capsid protein in vitro, Journal of Virology 66, 4874 (1992).

[4] S.-n. Kanesashi, K.-i. Ishizu, M.-a. Kawano, S.-i. Han, S. Tomita, H. Watanabe, K. Kataoka, and H. Handa, Simian virus 40 VP1 capsid protein forms polymorphic assemblies in vitro, Journal of General Virology 84, 1899 (2003).

[5] B. K. Ganser-Pornillos, U. K. von Schwedler, K. M. Stray, C. Aiken, and W. I. Sundquist, Assembly Properties of the Human Immunodeficiency Virus Type 1 CA Protein, Journal of Virology 78, 2545 (2004).

[6] J. Sun, C. DuFort, M.-C. Daniel, A. Murali, C. Chen, K. Gopinath, B. Stein, M. De, V. M. Rotello, A. Holzenburg, C. C. Kao, and B. Dragnea, Core-controlled polymorphism in virus-like particles, Proceedings of the National Academy of Sciences 104, 1354 (2007).

[7] H. D. Nguyen and C. L. I. Brooks, Generalized Structural Polymorphism in Self-Assembled Viral Particles, Nano Letters 8, 4574 (2008).

[8] L. Lavelle, M. Gingery, M. Phillips, W. M. Gelbart, C. M. Knobler, R. D. Cadena-Nava, J. R. Vega-Acosta, L. A. Pinedo-Torres, and J. Ruiz-Garcia, Phase Diagram of Self-assembled Viral Capsid Protein Polymorphs, The Journal of Physical Chemistry B 113, 3813 (2009).

[9] C. Butan, D. C. Winkler, J. B. Heymann, R. C. Craven, and A. C. Steven, RSV Capsid Polymorphism Correlates with Polymerization Efficiency and Envelope Glycoprotein Content: Implications that Nucleation Controls Morphogenesis, Journal of molecular biology 376, 1168 (2008).

[10] I. Seitz, S. Saarinen, E.-P. Kumpula, D. McNeale, E. Anaya-Plaza, V. Lampinen, V. P. Hytönen, F. Sains-bury, J. J. L. M. Cornelissen, V. Linko, J. T. Huisko-nen, and M. A. Kostiainen, DNA-origami-directed virus capsid polymorphism, Nature Nanotechnology 18, 1205 (2023).

[11] N. Harpell, A. L. Duran-Meza, H. Elmer, C. M. Kno-bler, J. A. Rodriguez, and W. M. Gelbart, Structural Polymorphs of Self-Assembled Brome Mosaic Virus Capsid Protein, The Journal of Physical Chemistry B 10.1021/acs.jpcb.5c08613 (2026).

[12] P. Kondylis, C. J. Schlicksup, N. E. Brunk, J. Zhou, A. Zlotnick, and S. C. Jacobson, Competition between Normative and Drug-Induced Virus Self-Assembly Ob-served with Single-Particle Methods, Journal of the American Chemical Society 141, 1251 (2019).

[13] K. Kra, S. Li, L. Gargowitsch, J. Degrouard, J. Pérez, R. Zandi, S. Bressanelli, and G. Tresset, Energetics and Kinetic Assembly Pathways of Hepatitis B Virus Capsids in the Presence of Antivirals, ACS Nano 17, 12723 (2023).

[14] C. A. Lutomski, N. A. Lyktey, E. E. Pierson, Z. Zhao, A. Zlotnick, and M. F. Jarrold, Multiple Pathways in Capsid Assembly, Journal of the American Chemical Society 140, 5784 (2018).

[15] R. Asor, L. Selzer, C. J. Schlicksup, Z. Zhao, A. Zlotnick, and U. Raviv, Assembly Reactions of Hepatitis B Capsid Protein into Capsid Nanoparticles Follow a Narrow Path through a Complex Reaction Landscape, ACS Nano 13, 7610 (2019).

[16] S. P. E. Timmermans, A. Ramezani, T. Montalvo, M. Nguyen, P. van der Schoot, J. C. M. van Hest, and R. Zandi, The Dynamics of Viruslike Capsid Assembly and Disassembly, Journal of the American Chemical Society 144, 12608 (2022).

[17] S. Li, G. Tresset, and R. Zandi, From disorder to icosahedral symmetry: How conformation-switching sub-units enable RNA virus assembly, Science Advances 11, eady7241 (2025).

[18] S. Li, P. Roy, A. Travesset, and R. Zandi, Why large icosahedral viruses need scaffolding proteins, Proceedings of the National Academy of Sciences 115, 10971 (2018).

[19] S. Panahandeh, S. Li, B. Dragnea, and R. Zandi, Virus Assembly Pathways Inside a Host Cell, ACS Nano 16, 317 (2022).

[20] D. L. D. Caspar and A. Klug, Physical Principles in the Construction of Regular Viruses, Cold Spring Harbor Symposia on Quantitative Biology 27, 1 (1962).

[21] J. E. Johnson and J. A. Speir, Quasi-equivalent viruses: A paradigm for protein assemblies1, Journal of Molecular Biology 269, 665 (1997).

[22] J. D. Perlmutter and M. F. Hagan, Mechanisms of Virus Assembly, Annual Review of Physical Chemistry 66, 217 (2015).

[23] Global Hepatitis Report 2024: Action for Access in Low- and Middle-Income Countries, 1st ed. (World Health Organization, Geneva, 2024).

[24] V. Soriano, P. Barreiro, E. Cachay, S. Kottilil, J. V. Fernandez-Montero, and C. de Mendoza, Advances in hepatitis B therapeutics, Therapeutic Advances in Infectious Disease 7, 2049936120965027 (2020).

[25] B. Nijampatnam and D. C. Liotta, Recent advances in the development of HBV capsid assembly modulators, Current Opinion in Chemical Biology Next Generation Therapeutics, 50, 73 (2019).

[26] V. Taverniti, G. Ligat, Y. Debing, D. B. Kum, T. F. Baumert, and E. R. Verrier, Capsid Assembly Modula-tors as Antiviral Agents against HBV: Molecular Mechanisms and Clinical Perspectives, Journal of Clinical Medicine 11, 1349 (2022).

[27] W. E. Delaney, R. Edwards, D. Colledge, T. Shaw, P. Furman, G. Painter, and S. Locarnini, Phenylprope-namide Derivatives AT-61 and AT-130 Inhibit Replication of Wild-Type and Lamivudine-Resistant Strains of Hepatitis B Virus In Vitro, Antimicrobial Agents and Chemotherapy 46, 3057 (2002).

[28] J. J. Feld, D. Colledge, V. Sozzi, R. Edwards, M. Lit-tlejohn, and S. A. Locarnini, The phenylpropenamide derivative AT-130 blocks HBV replication at the level of viral RNA packaging, Antiviral Research 76, 168 (2007).

[29] S. P. Katen, S. R. Chirapu, M. G. Finn, and A. Zlotnick, Trapping of Hepatitis B Virus Capsid Assembly Inter-mediates by Phenylpropenamide Assembly Accelerators, ACS Chemical Biology 5, 1125 (2010).

[30] S. J. Stray, C. R. Bourne, S. Punna, W. G. Lewis, M. G. Finn, and A. Zlotnick, A heteroaryldihydropyrimidine activates and can misdirect hepatitis B virus capsid assembly, Proceedings of the National Academy of Sciences 102, 8138 (2005).

[31] C. Bourne, S. Lee, B. Venkataiah, A. Lee, B. Korba, M. G. Finn, and A. Zlotnick, Small-Molecule Effectors of Hepatitis B Virus Capsid Assembly Give Insight into Virus Life Cycle, Journal of Virology 82, 10262 (2008).

[32] R. A. Crowther, N. A. Kiselev, B. Böttcher, J. A. Berri-man, G. P. Borisova, V. Ose, and P. Pumpens, Three-dimensional structure of hepatitis B virus core particles determined by electron cryomicroscopy, Cell 77, 943 (1994).

[33] P. T. Wingfield, S. J. Stahl, R. W. Williams, and A. C. Steven, Hepatitis Core Antigen Produced in Escherichia coli: Subunit Composition, Conformation Analysis, and in Vitro Capsid Assembly, Biochemistry 34, 4919 (1995).

[34] A. Zlotnick, N. Cheng, J. F. Conway, F. P. Booy, A. C. Steven, S. J. Stahl, and P. T. Wingfield, Dimorphism of Hepatitis B Virus Capsids Is Strongly Influenced by the C-Terminus of the Capsid Protein, Biochemistry 35, 7412 (1996).

[35] F. Mohajerani, B. Tyukodi, C. J. Schlicksup, J. A. Hadden-Perilla, A. Zlotnick, and M. F. Hagan, Multiscale Modeling of Hepatitis B Virus Capsid Assembly and Its Dimorphism, ACS Nano 16, 13845 (2022).

[36] C. J. Schlicksup, J. C.-Y. Wang, S. Francis, B. Venkatakrishnan, W. W. Turner, M. VanNieuwenhze, and A. Zlotnick, Hepatitis B virus core protein allosteric modulators can distort and disrupt intact capsids, eLife 7, e31473 (2018).

[37] J. A. Hadden, J. R. Perilla, C. J. Schlicksup, B. Venkatakrishnan, A. Zlotnick, and K. Schulten, All-atom molecular dynamics of the HBV capsid reveals in-sights into biological function and cryo-EM resolution limits, eLife 7, e32478 (2018).

[38] C. Pérez-Segura, B. C. Goh, and J. A. Hadden-Perilla, All-Atom MD Simulations of the HBV Capsid Complexed with AT130 Reveal Secondary and Tertiary Structural Changes and Mechanisms of Allostery, Viruses 13, 564 (2021).

[39] C. Pérez-Segura, B. C. Goh, and J. A. Hadden-Perilla, Mechanistic insights into CAM-induced disruption of HBV capsids revealed by all-atom MD simulations, PLOS Pathogens 22, e1013566 (2026).

[40] L. W. Scott, C.-P. Segura, J. A. Hadden-Perilla, and A. Zlotnick, Assembly-active and -inactive forms of HBV capsid protein provide distinctly different binding sites for capsid assembly modulators, Antimicrobial Agents and Chemotherapeutics (Submitted).

[41] P. Moerman, P. Van Der Schoot, and W. Kegel, Kinetics versus Thermodynamics in Virus Capsid Polymorphism, The Journal of Physical Chemistry B 120, 6003 (2016).

[42] K. Yang, J. M. Gonzalez, A. Ramezani, P. van der Schoot, and R. Zandi, Thermodynamic stability and kinetic control of capsid morphologies in hepatitis B virus, The Journal of Chemical Physics 164, 014112 (2026).

[43] G. M. Rotskoff and P. L. Geissler, Robust nonequilibrium pathways to microcompartment assembly, Proceedings of the National Academy of Sciences 115, 6341 (2018).

[44] P. Ceres and A. Zlotnick, Weak Protein-Protein Inter-actions Are Sufficient To Drive Assembly of Hepatitis B Virus Capsids, Biochemistry 41, 11525 (2002).

[45] R. Asor, C. J. Schlicksup, Z. Zhao, A. Zlotnick, and U. Raviv, Rapidly Forming Early Intermediate Structures Dictate the Pathway of Capsid Assembly, Journal of the American Chemical Society 142, 7868 (2020).

[46] S. Whitelam, E. H. Feng, M. F. Hagan, and P. L. Geissler, The role of collective motion in examples of coarsening and self-assembly, Soft Matter 5, 1251 (2009).

[47] M. F. Hagan and D. Chandler, Dynamic Pathways for Viral Capsid Assembly, Biophysical Journal 91, 42 (2006).

[48] J. Zhou, P. Kondylis, D. G. Haywood, Z. D. Harms, L. S. Lee, A. Zlotnick, and S. C. Jacobson, Characterization of Virus Capsids and Their Assembly Intermediates by Multicycle Resistive-Pulse Sensing with Four Pores in Series, Analytical Chemistry 90, 7267 (2018).

[49] P. Kondylis, C. J. Schlicksup, A. Zlotnick, and S. C. Ja-cobson, Analytical Techniques to Characterize the Structure, Properties, and Assembly of Virus Capsids, Analytical Chemistry 91, 622 (2019).

[50] J. Zhou, A. Zlotnick, and S. C. Jacobson, Disassembly of Single Virus Capsids Monitored in Real Time with Multicycle Resistive-Pulse Sensing, Analytical Chemistry 94, 985 (2022).

[51] S. M. Klein, A. Patterson, K. Young, M. P. Biever, L. M. Miller, A. Zlotnick, S. C. Jacobson, and M. F. Jarrold, Speed Matters: Directed Assembly of Icosahedral HPV Virus-Like Particles, Journal of the American Chemical Society 147, 24950 (2025).

[52] J.-H. Prinz, B. Keller, and F. Noé, Probing molecular kinetics with Markov models: Metastable states, transition pathways and spectroscopic observables, Physical Chemistry Chemical Physics 13, 16912 (2011).

[53] M. R. Perkett and M. F. Hagan, Using Markov state models to study self-assembly, The Journal of Chemical Physics 140, 214101 (2014).

[54] J. Ortega-Arroyo and P. Kukura, Interferometric scattering microscopy (iSCAT): New frontiers in ultrafast and ultrasensitive optical microscopy, Physical Chemistry Chemical Physics 14, 15625 (2012).

[55] G. Young, N. Hundt, D. Cole, A. Fineberg, J. Andrecka, A. Tyler, A. Olerinyova, A. Ansari, E. G. Marklund, M. P. Collier, S. A. Chandler, O. Tkachenko, J. Allen, M. Crispin, N. Billington, Y. Takagi, J. R. Sellers, C. Eichmann, P. Selenko, L. Frey, R. Riek, M. R. Galpin, W. B. Struwe, J. L. P. Benesch, and P. Kukura, Quantitative mass imaging of single biological macromolecules, Science 360, 423 (2018).

[56] R. F. Garmann, A. M. Goldfain, and V. N. Manoharan, Measurements of the self-assembly kinetics of individual viral capsids around their RNA genome, Proceedings of the National Academy of Sciences 116, 22485 (2019).

[57] R. F. Garmann, A. M. Goldfain, C. R. Tanimoto, C. E. Beren, F. F. Vasquez, D. A. Villarreal, C. M. Knobler, W. M. Gelbart, and V. N. Manoharan, Single-particle studies of the effects of RNA–protein interactions on the self-assembly of RNA virus particles, Proceedings of the National Academy of Sciences 119, e2206292119 (2022).

[58] E. D. B. Foley, M. S. Kushwah, G. Young, and P. Kukura, Mass photometry enables label-free tracking and mass measurement of single proteins on lipid bilayers, Nature Methods 18, 1247 (2021).

[59] R. Asor, D. Loewenthal, R. van Wee, J. L. P. Benesch, and P. Kukura, Mass Photometry, Annual Review of Bio-physics 54, 379 (2025).

[60] R. Asor, D. Loewenthal, D. Melnyk, T. K. Tan, and P. Kukura, Molecular-level observation of the self-assembly of a virus-like particle (2026).

[61] C. J. Valkner, N. Gibes, M. S. Xie, S. Kumar, S. Francis, B. Venkatakrishnan, I. Tsvetkova, A. Patterson, S. Nair, M. VanNieuwenhze, B. Dragnea, J. C.-Y. Wang, and A. Zlotnick, Negative cooperativity drives activity of capsid-directed antivirals against Hepatitis B Virus, Science Advances (In press).

[62] F. Amblard, Z. Chen, K. Kra, D. Patel, L. Bassit, L. Gargowitsch, J. Degrouard, V. rat, A.-A. Arteni, L. Matthews, S. M. Ravichandran, A. Jiang, A. Athanasiadis, G. Dangas, M. Abate, G. Tresset, J. Pronost, M. Salman, K. Verma, K. A. Kirby, E. Michailidis, H. de Rocquigny, D. Durantel, S. Bres-sanelli, S. G. Sarafianos, and R. F. Schinazi, Novel Dimeric Capsid Assembly Modulators as a Unique Class of Highly Potent Anti-HBV Agents, Journal of Medicinal Chemistry 68, 24196 (2025).

[63] L. Hozáková, B. Vokatá, T. Ruml, and P. Ulbrich, Targeting the Virus Capsid as a Tool to Fight RNA Viruses, Viruses 14, 174 (2022).

[64] A. J. Pak, J. M. A. Grime, A. Yu, and G. A. Voth, Off-Pathway Assembly: A Broad-Spectrum Mechanism of Action for Drugs That Undermine Controlled HIV-1 Viral Capsid Formation, Journal of the American Chemical Society 141, 10214 (2019).

[65] S. Segal-Maurer, E. DeJesus, H.-J. Stellbrink, A. Castagna, G. J. Richmond, G. I. Sinclair, K. Siripas-sorn, P. J. Ruane, M. Berhe, H. Wang, N. A. Margot, H. Dvory-Sobol, R. H. Hyland, D. M. Brainard, M. S. Rhee, J. M. Baeten, and J.-M. Molina, Capsid Inhibition with Lenacapavir in Multidrug-Resistant HIV-1 Infection, New England Journal of Medicine 386, 1793 (2022).

[66] N. F. B. dos Santos, J. A. Lewis, M. Hansen, M. J. B. Pereira, D. E. Christensen, W. I. Sundquist, B. K. Ganser-Pornillos, and O. Pornillos, Lenacapavir allosterically remodels the HIV-1 capsid (2026).

[67] M. Gupta, C. Waltmann, N. Renner, Y. Wang, L. C. James, D. A. Jacques, T. Böcking, and G. A. Voth, Mechanistic insights into lenacapavir-induced off-pathway HIV-1 capsid assembly, Proceedings of the National Academy of Sciences 123, e2524995123 (2026).

[68] R. Arai, Hierarchical design of artificial proteins and complexes toward synthetic structural biology, Biophysical Reviews 10, 391 (2018).

[69] P. W. K. Rothemund, Folding DNA to create nanoscale shapes and patterns, Nature 440, 297 (2006).

[70] S. M. Douglas, H. Dietz, T. Liedl, B. Högberg, F. Graf, and W. M. Shih, Self-assembly of DNA into nanoscale three-dimensional shapes, Nature 459, 414 (2009).

[71] T. Gerling, K. F. Wagenbauer, A. M. Neuner, and H. Di-etz, Dynamic DNA devices and assemblies formed by shape-complementary, non–base pairing 3D components, Science 347, 1446 (2015).

[72] W. M. Jacobs and W. B. Rogers, Assembly of Complex Colloidal Systems Using DNA, Annual Review of Condensed Matter Physics 16, 443 (2025).

[73] J. E. Padilla, C. Colovos, and T. O. Yeates, Nanohe-dra: Using symmetry to design self assembling protein cages, layers, crystals, and filaments, Proceedings of the National Academy of Sciences 98, 2217 (2001).

[74] Y.-T. Lai, D. Cascio, and T. O. Yeates, Structure of a 16-nm Cage Designed by Using Protein Oligomers, Science 336, 1129 (2012).

[75] P.-S. Huang, S. E. Boyken, and D. Baker, The coming of age of de novo protein design, Nature 537, 320 (2016).

[76] Z. Li, S. Wang, U. Nattermann, A. K. Bera, A. J. Borst, M. Y. Yaman, M. J. Bick, E. C. Yang, W. Sheffler, B. Lee, S. Seifert, G. L. Hura, H. Nguyen, A. Kang, R. Dalal, J. M. Lubner, Y. Hsia, H. Haddox, A. Courbet, Q. Dowling, M. Miranda, A. Favor, A. Etemadi, N. I. Edman, W. Yang, C. Weidle, B. Sankaran, B. Negahdari, M. B. Ross, D. S. Ginger, and D. Baker, Accurate computational design of three-dimensional protein crystals, Nature Materials 22, 1556 (2023).

[77] T. Kortemme, De novo protein design—From new structures to programmable functions, Cell 187, 526 (2024).

[78] S. Wang, A. Favor, R. D. Kibler, J. M. Lubner, A. J. Borst, N. Coudray, R. L. Redler, H. T. Chiang, W. Shef-fler, Y. Hsia, N. P. Bethel, Z. Li, D. C. Ekiert, G. Bhabha, L. D. Pozzo, and D. Baker, Bond-centric modular design of protein assemblies, Nature Materials 24, 1644 (2025).

[79] W. Yang, S. Wang, G. R. Lee, J. Z. Zhang, A. Courbet, D. Juergens, X. Wang, T. Schlichthaerle, M. Abedi, R. Ragotte, L. An, I. Kalvet, S. Pellock, L. Mihal-jevic, C. Glasscock, A. Pillai, A. Broerman, N. En-nist, E. Haefner, N. McNamara-Bordewick, I. Haydon, L. Stewart, G. Bhardwaj, and D. Baker, The past, present and future of de novo protein design, Nature 652, 1139 (2026).

[80] D. Y. Zhang and G. Seelig, Dynamic DNA nanotechnology using strand-displacement reactions, Nature Chemistry 3, 103 (2011).

[81] R. A. Langan, S. E. Boyken, A. H. Ng, J. A. Samson, G. Dods, A. M. Westbrook, T. H. Nguyen, M. J. La-joie, Z. Chen, S. Berger, V. K. Mulligan, J. E. Due-ber, W. R. P. Novak, H. El-Samad, and D. Baker, De novo design of bioactive protein switches, Nature 572, 205 (2019).

[82] A. Pillai, A. Idris, A. Philomin, C. Weidle, R. Skotheim, P. J. Y. Leung, A. Broerman, C. Demakis, A. J. Borst, F. Praetorius, and D. Baker, De novo design of allosterically switchable protein assemblies, Nature 632, 911 (2024).

[83] T. E. Videbæk, D. Hayakawa, G. M. Grason, M. F. Ha-gan, S. Fraden, and W. B. Rogers, Economical routes to size-specific assembly of self-closing structures, Science Advances 10, eado5979 (2024).

[84] E. Krissinel and K. Henrick, Inference of macromolecular assemblies from crystalline state, Journal of Molecular Biology 372, 774 (2007).

[85] ‘Protein interfaces, surfaces and assemblies’ service PISA at the European Bioinformatics Institute. (http://www.ebi.ac.uk/pdbe/protint/pistart.html)., E. Krissinel, and K. Henrick, Inference of Macromolecular Assemblies from Crystalline State, Journal of Molecular Biology 372, 774 (2007).

[86] M. F. Hagan and O. M. Elrad, Understanding the Concentration Dependence of Viral Capsid Assembly Kinetics—the Origin of the Lag Time and Identifying the Critical Nucleus Size, Biophysical Journal 98, 1065 (2010).

[87] L. Selzer, S. Katen, and A. Zlotnick, A Disulfide in HBV Core Protein Dimer Allosterically Modifies Capsid Assembly and Stability, Biophysical Journal 106, 60a(2014).

[88] W. Humphrey, A. Dalke, and K. Schulten, VMD: Visual molecular dynamics, Journal of Molecular Graphics 14, 33 (1996).

## References

[1] F. Mohajerani, B. Tyukodi, C. J. Schlicksup, J. A. Hadden-Perilla, A. Zlotnick, and M. F. Hagan, Multiscale Modeling of Hepatitis B Virus Capsid Assembly and Its Dimorphism, ACS Nano 16, 13845 (2022).

[2] L. Kettner, Designing a data structure for polyhedral surfaces, in Proceedings of the Fourteenth Annual Symposium on Computational Geometry -SCG ‘98 (ACM Press, Minneapolis, Minnesota, United States, 1998) pp. 146–154.

[3] D. E. Muller and F. P. Preparata, Finding the intersection of two convex polyhedra, Theoretical Computer Science 7, 217 (1978).

[4] S. Li, R. Zandi, A. Travesset, and G. M. Grason, Ground States of Crystalline Caps: Generalized Jellium on Curved Space, Physical Review Letters 123, 145501 (2019).

[5] G. M. Rotskoff and P. L. Geissler, Robust nonequilibrium pathways to microcompartment assembly, Proceedings of the National Academy of Sciences 115, 6341 (2018).

[6] W. Wu, N. R. Watts, N. Cheng, R. Huang, A. C. Steven, and P. T. Wingfield, Expression of quasi-equivalence and capsid dimorphism in the Hepadnaviridae, PLOS Computational Biology 16, e1007782 (2020).

[7] R. Asor, L. Selzer, C. J. Schlicksup, Z. Zhao, A. Zlotnick, and U. Raviv, Assembly Reactions of Hepatitis B Capsid Protein into Capsid Nanoparticles Follow a Narrow Path through a Complex Reaction Landscape, ACS Nano 13, 7610 (2019).

[8] P. Buzón, S. Maity, P. Christodoulis, M. J. Wiertsema, S. Dunkelbarger, C. Kim, G. J. Wuite, A. Zlotnick, and W. H. Roos, Virus self-assembly proceeds through contact-rich energy minima, Science Advances 7, eabg0811 (2021).

[9] R. C. Oliver, W. Potrzebowski, S. M. Najibi, M. N. Pedersen, L. Arleth, N. Mahmoudi, and I. André, Assembly of Capsids from Hepatitis B Virus Core Protein Progresses through Highly Populated Intermediates in the Presence and Absence of RNA, ACS Nano 14, 10226 (2020).

[10] N. Metropolis, A. W. Rosenbluth, M. N. Rosenbluth, A. H. Teller, and E. Teller, Equation of State Calculations by Fast Computing Machines, The Journal of Chemical Physics 21, 1087 (1953).

[11] W. K. Hastings, Monte Carlo sampling methods using Markov chains and their applications, Biometrika 57, 97 (1970).

[12] M. F. Hagan and D. Chandler, Dynamic Pathways for Viral Capsid Assembly, Biophysical Journal 91, 42 (2006).

[13] T. Zhang and R. Schwartz, Simulation Study of the Contribution of Oligomer/Oligomer Binding to Capsid Assembly Kinetics, Biophysical Journal 90, 57 (2006).

[14] W. C. Swope, J. W. Pitera, F. Suits, M. Pitman, M. Eleftheriou, B. G. Fitch, R. S. Germain, A. Rayshubski, T. J. C. Ward, Y. Zhestkov, and R. Zhou, Describing Protein Folding Kinetics by Molecular Dynamics Simulations. 2. Example Applications to Alanine Dipeptide and a β-Hairpin Peptide, The Journal of Physical Chemistry B 108, 6582 (2004).

